# Itch receptor MRGPRX4 interacts with the receptor activity-modifying proteins (RAMPs)

**DOI:** 10.1101/2022.12.06.519316

**Authors:** Ilana B. Kotliar, Emilie Ceraudo, Kevin Kemelmakher-Liben, Deena A. Oren, Emily Lorenzen, Tea Dodig-Crnković, Mizuho Horioka-Duplix, Thomas Huber, Jochen M. Schwenk, Thomas P. Sakmar

**Affiliations:** Laboratory of Chemical Biology and Signal Transduction, The Rockefeller University, 1230 York Ave, New York, NY 10065, USA; Tri-Institutional PhD Program in Chemical Biology, New York, NY 10065, USA; Structural Biology Resource Center, The Rockefeller University, 1230 York Ave, New York, NY 10065, USA; Science for Life Laboratory, Department of Protein Science, School of Engineering Sciences in Chemistry, Biotechnology and Health, KTH Royal Institute of Technology, 171 65 Solna, Sweden; Department of Neurobiology, Care Sciences and Society, Section for Neurogeriatrics, Karolinska Institutet, 171 64 Solna, Sweden

## Abstract

Cholestatic itch is a severe and debilitating symptom in liver diseases with limited treatment options. The class A G protein-coupled receptor (GPCR) Mas-related GPCR subtype X4 (MRGPRX4) has been identified as a receptor for bile acids, which are potential cholestatic pruritogens. An increasing number of GPCRs have been shown to interact with receptor activity-modifying proteins (RAMPs), which can modulate different aspects of GPCR biology. Using a combination of multiplexed immunoassay and proximity ligation assay we show that MRGPRX4 interacts with RAMPs. The interaction of MRGPRX4 with RAMP2, but not RAMP1 or 3, causes attenuation of basal and agonist-dependent signaling, which correlates with a decrease of MRGPRX4 cell surface expression as measured using a quantitative NanoBRET pulse-chase assay. Finally, we use AlphaFold Multimer to predict the structure of the MRGPRX4-RAMP2 complex. The discovery that RAMP2 regulates MRGPRX4 may have direct implications for future drug development for cholestatic itch.

## Introduction

Cholestatic itch, or pruritus, is a severe and potentially debilitating symptom that affects more than 80% of patients with cholestatic liver diseases, including primary biliary cholangitis and end stage liver cirrhosis^1^. Cholestasis is generally associated with increased plasma levels of bile acids (BA) and bilirubin. The G protein-coupled receptor (GPCR) called Mas-related GPCR subtype X4 (MRGPRX4) has recently been deorphanized as a receptor for BAs and bilirubin^2–4^. The activation of MRGPRX4 contributes to BA- and bilirubin-induced itch in transgenic mice^2, 3^. The activation of MRGPRX4 triggers itch sensation in human subjects and elevated levels of BAs found in cholestatic itch patients are sufficient to activate MRGPRX4^4^. Together these reports suggest that MRGPRX4 mediates itch in response to BA and bilirubin^1–4^.

MRGPRX4 is a class A, delta subfamily GPCR expressed primarily in small-diameter sensory neurons of the dorsal root ganglia (DRG) and trigeminal ganglia (TG)^5–, 8^ and in skin keratinocytes^9^. It is reported to couple with Gq signaling pathways to activate phospholipase C (PLC) β to generate the second messenger inositol 1,4,5-trisphosphate (IP3), which mediates intracellular calcium Ca^2+^ release prior to being degraded into inositol monophosphate (IP1)^2–4^. MRGPRX4 is a potential target for drug development efforts to treat cholestatic itch associated with liver diseases. In addition, certain drugs such as nateglinide, a potassium ATP channel blocker used for treatment of type 2 diabetes, are hypothesized to cause pruritus and urticarial rash by activating MRGPRX4 as an off-target side effect^4, 10^.

BA metabolism is complex. The primary BAs cholic acid (CA) and chenodeoxycholic acid (CDCA) are secreted into bile as glycine or taurine conjugates and metabolized by gut bacteria into the secondary BAs deoxycholic acid (DCA), ursodeoxycholic acid (UDCA), and lithocholic acid (LCA)^11, 12^. Taurodeoxycholic acid (TDCA) is a conjugated form of DCA that is also present in bile^11, 12^. In humans, the circulating pool of BAs that can reach pharmacological levels in serum consists mainly of CA, CDCA, and DCA^11^. A systematic study of the pharmacology of BAs at MRGPRX4 and the potential role of GPCR accessory proteins in the regulation of MRGPRX4 has not been carried out, which are both required for drug discovery efforts.

Here, we show that MRGPRX4 interacts with a receptor activity-modifying protein (RAMP) that affects its ability to signal in response to treatment with BAs. We investigated the activation of downstream Gq signaling pathways and the β-arrestin recruitment to MRGPRX4 to assess the effects of each of the three RAMPs. We observed that although RAMP2 and RAMP3 interact with MRGPRX4, only RAMP2 modulates signaling by down-regulating receptor cell surface expression and total expression. Furthermore, we identified that among the BAs studied, DCA is a biased agonist and mediates Gq signaling preferentially to β-arrestin recruitment at MRGPRX4. In addition, we employed AlphaFold Multimer to generate the predicted structure of the MRGPRX4-RAMP2 complex. These results provide important information about the biology and pharmacology of MRGPRX4, an important potential drug target.

## Results

### MRGPRX4 signals through Gq and displays high IP1 basal activity

We employed the homogenous time-resolved fluorescence (HTRF) IP1 accumulation assay to characterize MRGPRX4 Gq protein signaling. Dose-response curves for the known MRGPRX4 agonists DCA, TDCA, and UDCA were compared with that of the diabetes drug nateglinide (**Fig. 1a**, **Supplementary Fig. 1a**)^2, 10^. Nateglinide displayed higher potency and higher efficacy than any of agonist BAs tested (**Supplementary Table 1**). For example, the EC_50_ concentration for nateglinide (10.6 µM) was approximately two-fold lower than that of DCA (19.2 µM) and more than five-fold lower than TDCA or UDCA.

**Fig. 1.**
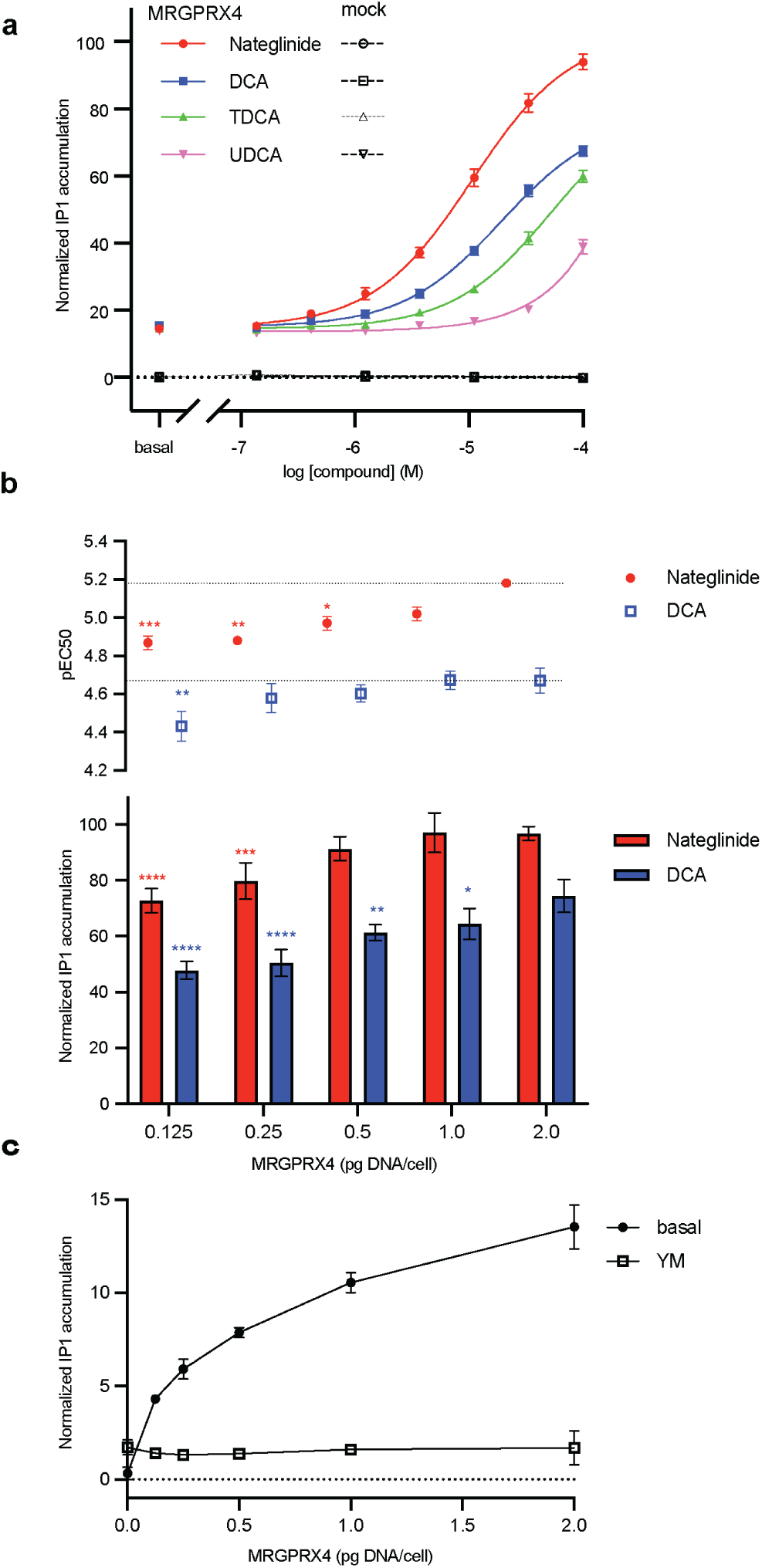
Agonist dose-response curves and determination of basal signaling. **a** MRGPRX4 agonist dose-dependent IP1 accumulation for nateglinide (red circles), DCA (blue squares), TDCA (green triangles), and UDCA (pink reverse triangles). Fitting parameters are provided in **Supplementary Table 1**. **b** IP1 accumulation induced by nateglinide (red) and DCA (blue) in cells expressing different amounts of MRGPRX4. The pEC50 plot displays the midpoint of the dose-response curves in **Supplementary Fig. 1**. The bar graphs display agonist-dependent activity as the difference between the normalized basal and maximal IP1 accumulation values for the dose-response curves in **Supplementary Fig. 1**. Data are expressed as the mean of the normalized IP1 accumulation. The error bars represent the standard errors of the mean (SEM). The presented data are from three independent experiments performed in four technical replicates. IP1 accumulation is normalized to 100 µM nateglinide-stimulated MRGPRX4. Statistical significance was determined by ordinary one-way ANOVA followed by Dunnett’s multiple comparisons test to MRGPRX4 (2 pg DNA/cell). Statistical significance, \*\*\*\**p* < 0.0001, \*\*\**p* < 0.001, \*\**p* < 0.01, \**p* < 0.05 with red * for nateglinide and blue * for DCA comparisons (see **Supplementary Table 2**). **c** Basal IP1 accumulation in the absence of agonist in cells expressing different amounts of MRGPRX4 with or without the Gq inhibitor YM254890 (YM).

Therefore, for subsequent characterization of MRGPRX4 we focused primarily on stimulation of the receptor with nateglinide and DCA. Both induced IP1 accumulation that scaled proportionally with the level of MRGPRX4 expression (**Fig. 1b**, **Supplementary Fig. 1b-c**, **Supplementary Table 1**). Additionally, the potency of both agonists, as reflected by the pEC_50_ values, increased with the level of MRGPRX4 expression (**Fig. 1b**). Nateglinide activated MRGPRX4 more potently than DCA at all expression levels. The nateglinide and DCA responses were inhibited by the Gq inhibitor YM254890 (YM) and the phospholipase C (PLC) inhibitor U73122, confirming that both agonists signal through Gq coupling to MRGPRX4 and activate the PLC pathway (**Supplementary Fig 1d-e**). Surprisingly, MRGPRX4 showed a high basal IP1 activity that positively scaled with the expression of MRGPRX4. The basal activity was abolished by YM and U73122 (**Fig. 1c, Supplementary Fig. 1e**). Together, these data show that MRGPRX4 signals through the Gq-PLC pathway in basal and agonist-dependent conditions.

### MRGPRX4 interacts with RAMPs

We employed a multiplexed suspension bead array (SBA) immunoassay to identify MRGPRX4-RAMP interactions. We tested for MRGPRX4-RAMP complexes derived from cells expressing dual epitope-tagged MRGPRX4 (HA-MRGPRX4-1D4) and complementary dual epitope-tagged RAMPs (FLAG-RAMP-OLLAS) (**Fig. 2a**)^13^. The multiplexed nature of the SBA assay allowed us to simultaneously validate the expression of MRGPRX4 and each RAMP with two different capture-detection schemes each (**Supplementary Fig. 2**). First, we showed highly significant expression of MRGPRX4 and all three RAMPs. Next, we subjected these samples to multiplexed analysis and applied eight different epitope tag-based capture-detection schemes to rank the RAMPs ability to interact with MRGPRX4 (**Fig. 2b, Supplementary Table 2**). MRGPRX4-RAMP2 and MRGPRX4-RAMP3 complexes were detected with high significance across all eight (100%) capture-detection schemes. MRGPRX4-RAMP1 complexes were only detected with high significance (*p* < 0.0001) by five of the eight (62.5%) approaches tested. The results across all schemes are summarized in **Fig. 2c**. Together, these data suggest that MRGPRX4 most probably forms complexes with RAMP2 and RAMP3.

**Fig. 2.**
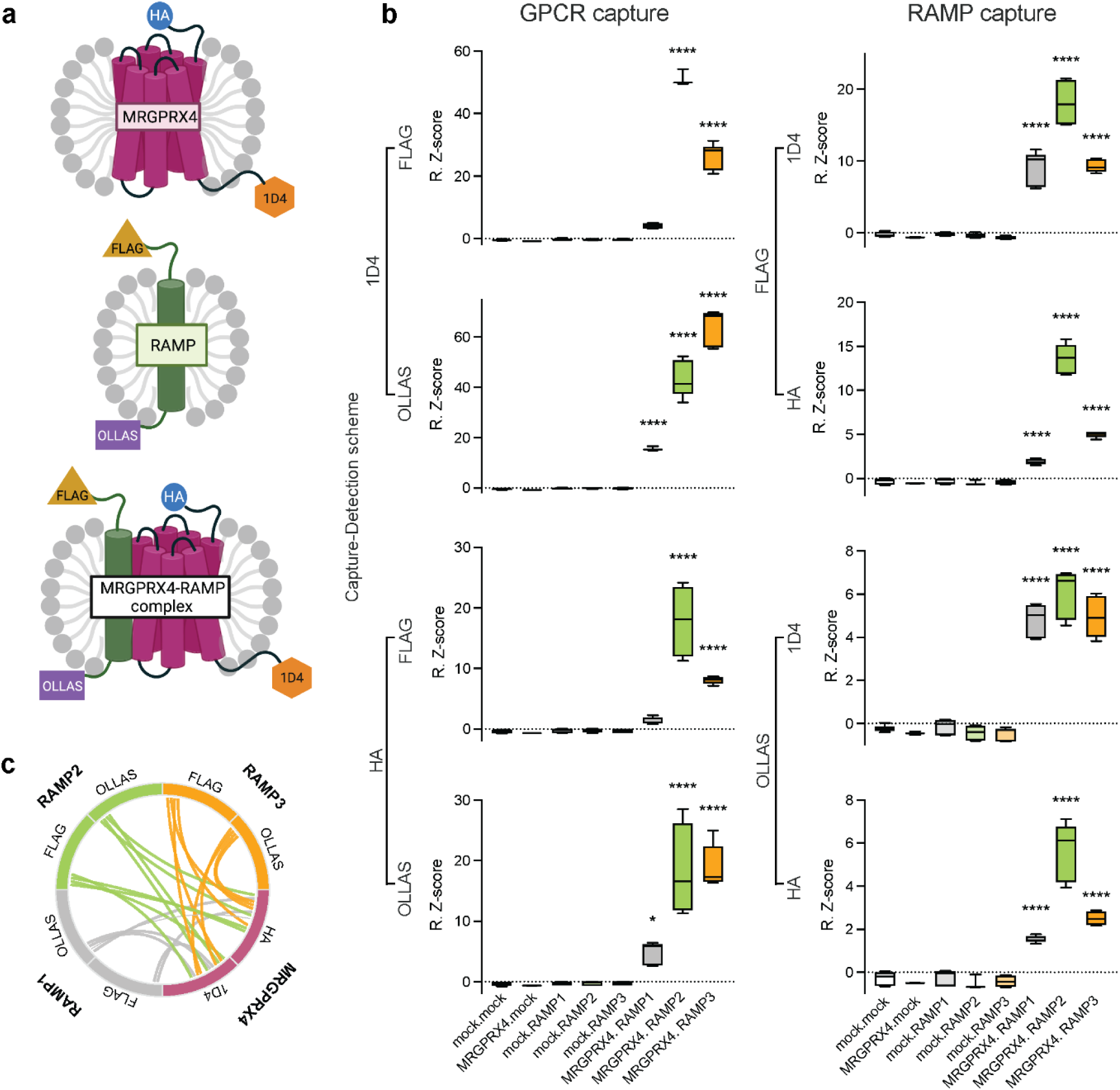
Discovery of MRGPRX4-RAMP complexes in solubilized cell membranes by suspension bead array (SBA) assay. **a** MRGPRX4 was tagged at its N-terminal and C-terminal tail with HA and 1D4 mAb epitopes, respectively. RAMPs 1-3 were tagged at their N-terminal and C-terminal tails with FLAG and OLLAS mAb epitopes, respectively^13^. **b** Lysates from Freestyle 293 cells transfected with epitope-tagged MRGPRX4, or each RAMP construct, or co-transfected pairwise with MRGPRX4 and each RAMP were incubated with the SBA, which included beads conjugated to mAbs against the four epitope tags. MRGPRX4, RAMPs, and MRGPRX4-RAMP complexes were captured on the beads in a multiplexed fashion. **b** The eight possible capture-detection schemes for the complexes are shown. (**b**, *left column*) MRGPRX4 was captured using anti-1D4 mAb or anti-HA mAb, and the MRGPRX4-RAMP complex was detected using PE-conjugated anti-FLAG mAb or PE-conjugated anti-OLLAS mAb. (**b**, *right column*) The RAMP was captured using anti-FLAG mAb or anti-OLLAS mAb, and the MRGPRX4-RAMP complex was detected using PE-conjugated anti-1D4 mAb or PE-conjugated anti-HA mAb. Sample names are listed at the bottom of each column using the format “transfected GPCR name (if any).transfected RAMP name (if any)” and the boxes are color coded. Data are plotted as Robust Z-scores (R.Z-scores) and represent measurements from three independent experiments performed in duplicate, except for MRGPRX4.mock data which is from one experiment performed in duplicate. The extremes of the box and whiskers plots represent the maximum and minimum values. The box is from the 25^th^ to the 75^th^ percentile. Statistical significance was determined by ordinary one-way ANOVA followed by Dunnett’s multiple comparisons test to mock.mock (see **Supplementary Table 2**) (\*\*\*\**p* < 0.0001, \**p* < 0.05, if not marked then not significant). **c** Graphical summary of MRGPRX4-RAMP interactions detected by SBA assay across all capture-detection schemes. Curved lines show pairwise MRGPRX4-RAMP interactions. The labels around the circumference indicate the capture-detection scheme. The statistical significance for each capture-detection pair is represented by the thickness of the curved lines. *p* ≤ 0.05 is given an arbitrary thickness of 1 and *p* < 0.0001 a thickness of 4. Color code: MRGPRX4 maroon, RAMP1 gray, RAMP2 lime, RAMP3 tangerine.

### Validation of MRGPRX4-RAMP complexes in cells

We employed the Proximity Ligation Assay (PLA) to detect the presence of MRGPRX4-RAMP complexes in cell membranes. We compared PLA puncta counts from cells expressing MRGPRX4 alone to those co-expressing MRGPRX4 with each of the three RAMPs (**Fig. 3, Supplementary Table 2**). The PLA puncta counts for MRGPRX4 co-expressed with either RAMP2 and RAMP3 reached high statistical significance (p<0.0001), providing additional evidence for MRGPRX4-RAMP2 and MRGPRX4-RAMP3 complex formation (**Fig. 3a-b, Supplementary Table 2**). The PLA puncta in cells co-expressing MGRPRX4 and RAMP2 appeared to be intracellular, compared to the largely cell surface-localized PLA puncta of cells co-expressing MRGPRX4 and RAMP3. We observed that the PLA puncta count for cells expressing MRGPRX4 and RAMP1 differed only slightly from that for cells expressing MRGPRX4 alone, although the interaction did reach statistical significance.

**Fig. 3.**
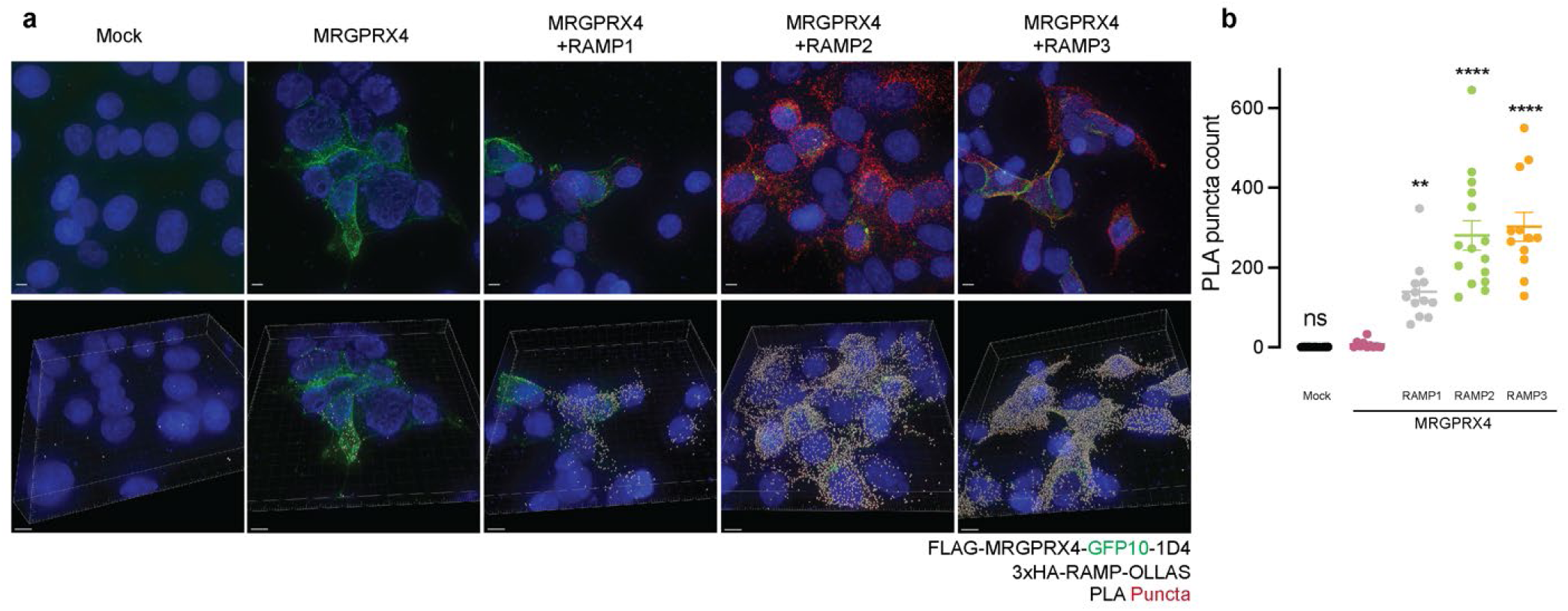
Validation of MRGPRX4-RAMP complex formation in cell membranes by proximity ligation assay (PLA). HEK293T cells were co-transfected with epitope-tagged MRGPRX4 and RAMPs and then incubated with anti-HA and anti-FLAG Abs. PLA was then carried out to quantitate MRGPRX4-RAMP interactions^13^. The number of PLA puncta per cell for each *Z*-stack captured was measured. Each *Z*-stack is of a different field of view. **a** Representative images of cells transfected with MRGPRX4 or MRGPRX4 with each RAMP subjected to PLA analysis. The top row shows the maximum projection of the *Z*-stack, which is the maximum signal intensity for each channel at each point across all slices. The bottom row shows snapshots from quantification of puncta performed in Imaris. Scale bars: 5 µm (top row images), 8 µm (bottom row images). DAPI blue, PLA puncta red, Imaris spots gray. **b** PLA puncta counts from Imaris quantitation of all PLA images collected. Data are from three independent experiments performed with triplicates for mock and four to five replicates for all other conditions. Data are shown as mean ± SEM with individual data points overlaid. Statistical significance was determined by an ordinary one-way ANOVA followed by Dunnett’s multiple comparisons test to MRGPRX4 alone (see **Supplementary Table 2**) (\*\*\*\**p* < 0.0001, **p < 0.01, ns, not significant).

Next, we studied the expression of a MRGPRX4 construct fused with GFP10. The fluorescence of the MRGPRX4-GFP10 fusion was largely localized at the cell surface when expressed alone. Similarly, MRGPRX4-GFP10 appeared to have plasma membrane localization when co-expressed with RAMP1 or RAMP3. Conversely, MRGPRX4-GFP10 was largely intracellular when co-expressed with RAMP2 (**Fig. 3a**). Together, the SBA assay and PLA data suggest that MRGPRX4 can most likely form stable complexes with RAMP2 and RAMP3. Further, the cellular localization of MRGPRX4 appears to be affected by co-expression with RAMP2, but not co-expression with RAMP1 or RAMP3.

### RAMP2 co-expression with MRGPRX4 alters its Gq signaling

To interrogate the functional consequences of the putatively identified MRGPRX4-RAMP interactions, we measured the effect of RAMP co-expression on MRGPRX4-dependent IP1 accumulation (**Fig. 4, Supplementary Fig. 3**). First, we measured IP1 accumulation mediated by the four different engineered MRGPRX4 constructs used in this study. We showed that the constructs all have comparable functionality (**Supplementary Fig. 4a-f**). We validated the functionality of the different epitope-tagged RAMP constructs by characterizing the IP1 accumulation promoted by the prototypical RAMP-interacting GPCR, calcitonin receptor-like receptor (CALCRL) (**Supplementary Fig. 4g-l**)^14^. We showed that RAMP expression is not affected by MRGPRX4 co-expression (**Supplementary Fig. 5m**). Next, we characterized agonist-dependent IP1 accumulation promoted by MRGPRX4 in the presence or absence of different levels of each RAMP. We showed that increasing levels of RAMP1 did not significantly alter the nateglinide- and DCA-induced IP1 accumulation mediated by MRGPRX4 (**Fig. 4a, Supplementary Fig. 3, Supplementary Table 1,2**). We observed a substantial attenuation of agonist-dependent response when MRGPRX4 was co-expressed with RAMP2, an effect that scaled with the amount of RAMP2 expressed.

**Fig. 4.**
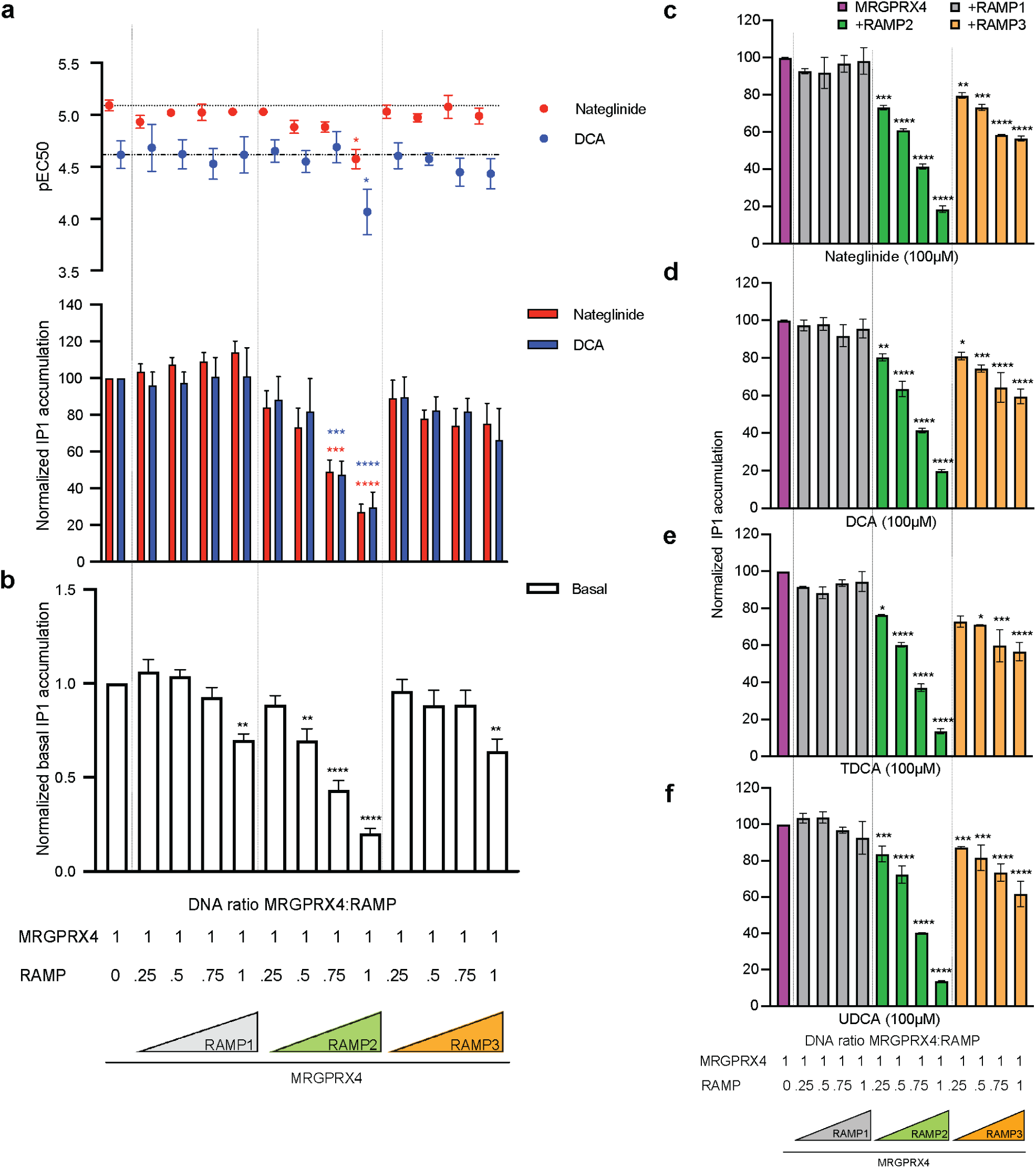
The effect of RAMP co-expression on MRGPRX4-mediated basal and agonist induced IP1 accumulation. **a** IP1 accumulation induced by nateglinide and DCA in cells expressing MRGPRX4 alone or with increasing amounts of each RAMP. (top) The data in the pEC50 plot represents the midpoint of the dose-response curve for each condition (nateglinide, red; DCA, blue). (bottom) The bar graphs represent agonist-dependent IP1 accumulation as the span between the endpoints of the dose-responses curves (see **Supplementary Figure 3**). Fitting parameters are provided in **Supplementary Table 1**. Data are expressed as the mean ± SEM of the normalized IP1 accumulation. Normalization is over the maximum of each agonist and the minimum of mock basal. The statistical significance was determined by ordinary one-way ANOVA followed by Dunnett’s multiple comparisons test to MRGPRX4 alone (see **Supplementary Table 2** for numerical parameters). **b** Basal IP1 accumulation in cells expressing MRGPRX4 alone or with increasing amounts of each RAMP. IP1 accumulation is normalized to MRGPRX4 basal and mock basal. **c-f** MRGPRX4-mediated IP1 accumulation in the presence of increasing expression of each RAMP following treatment with 100 µM **c** nateglinide, **d** DCA, **e** TDCA, or **f** UDCA. Data are expressed as the mean ± SEM. Statistical significance was determined by ordinary one-way ANOVA followed by Dunnett’s multiple comparisons test to MRGPRX4 basal (**b**) or MRGPRX4 stimulated with each agonist separately (**a**) (see **Supplementary Table 2**). (\*\*\*\**p* < 0.0001, \**p* < 0.05, if not marked then not significant). Data are from three independent experiments performed in four technical replicates (**a-b**) and two independent experiments performed in three technical replicates (**c-f**).

Interestingly, increasing levels of RAMP3 co-expression did not significantly alter IP1 accumulation. In all cases, the effect of the RAMPs was similar for both DCA and nateglinide-dependent activation. Subsequently, we investigated the effect of RAMP co-expression on the basal IP1 accumulation elicited by MRGPRX4 (**Fig. 4b, Supplementary Table 2**). We showed that RAMP1 and RAMP3 co-expression did not significantly alter MRGPRX4 basal signaling except at the highest level of RAMP expression. Conversely, RAMP2 co-expression resulted in a significant decrease of MRGPRX4 basal IP1 activity, an effect that scaled with the amount of RAMP2 expressed. To test whether the effect of RAMP co-expression with MRGPRX4 reflected agonist selectivity, we compared IP1 accumulation induced by the previously characterized agonists (**Fig. 4c-f, Supplementary Fig. 3, Supplementary Table 2**). RAMP2 and RAMP3, but not RAMP1, co-expression with MRGPRX4 correlated with decreased IP1 accumulation across all ligands compared with MRGPRX4 alone. The effect of MRGPRX4-RAMP2 co-expression was the most pronounced, with a maximal IP1 accumulation reduction of approximately 80% at the highest level of RAMP2 expression for all ligands. Taken together, these data demonstrate that co-expression of RAMP2, but not RAMP1 or RAMP3, with MRGPRX4 correlates to a strong decrease in MRGPRX4 Gq-mediated activation that is not agonist-selective. The observed trend of the effect of each RAMP is similar for basal and agonist-dependent signaling.

### MRGPRX4 differentially recruits β-arrestins with limited effect of RAMP co-expression

We used a bioluminescence resonance energy transfer two (BRET^2^) assay to characterize the β-arrestin1 and β-arrestin2 recruitment to MRGPRX4 in the presence or absence of each RAMP. First, we measured β-arrestin1 and β-arrestin2 recruitment to MRGPRX4 under basal and agonist-dependent conditions (**Fig. 5a,b**). MRGPRX4 did not recruit β-arrestin1 upon DCA stimulation and displayed very low β-arrestin1 recruitment upon nateglinide treatment (**Fig. 5a**). Conversely, MRGPRX4 recruited β-arrestin2 more strongly than β-arrestin1 in response to nateglinide, but not in response to DCA, an effect that scaled with the amount of MRGPRX4 expressed (**Fig. 5b**). Next, we studied the time-dependence of β-arrestin2 recruitment to MRGPRX4 (**Fig. 5c, Supplementary Table 3**). MRGPRX4 recruited β-arrestin2 quickly, with the peak of nateglinide-dependent recruitment to MRGPRX4 occurring at three minutes, followed by a reduction back to near baseline. On the other hand, we did not observe any β-arrestin2 recruitment in response to DCA, which is consistent with **Fig. 5b**. We used CALCRL, which requires RAMP co-expression to traffic to the cell membrane, co-expressed with RAMP2 as a positive control^14^. As a class B GPCR, CALCRL in complex with RAMP2 strongly recruits β-arrestin2 in response to adrenomedullin, while more weakly recruiting β-arrestin1, as shown in **Fig. 5d**^15^.

**Fig. 5.**
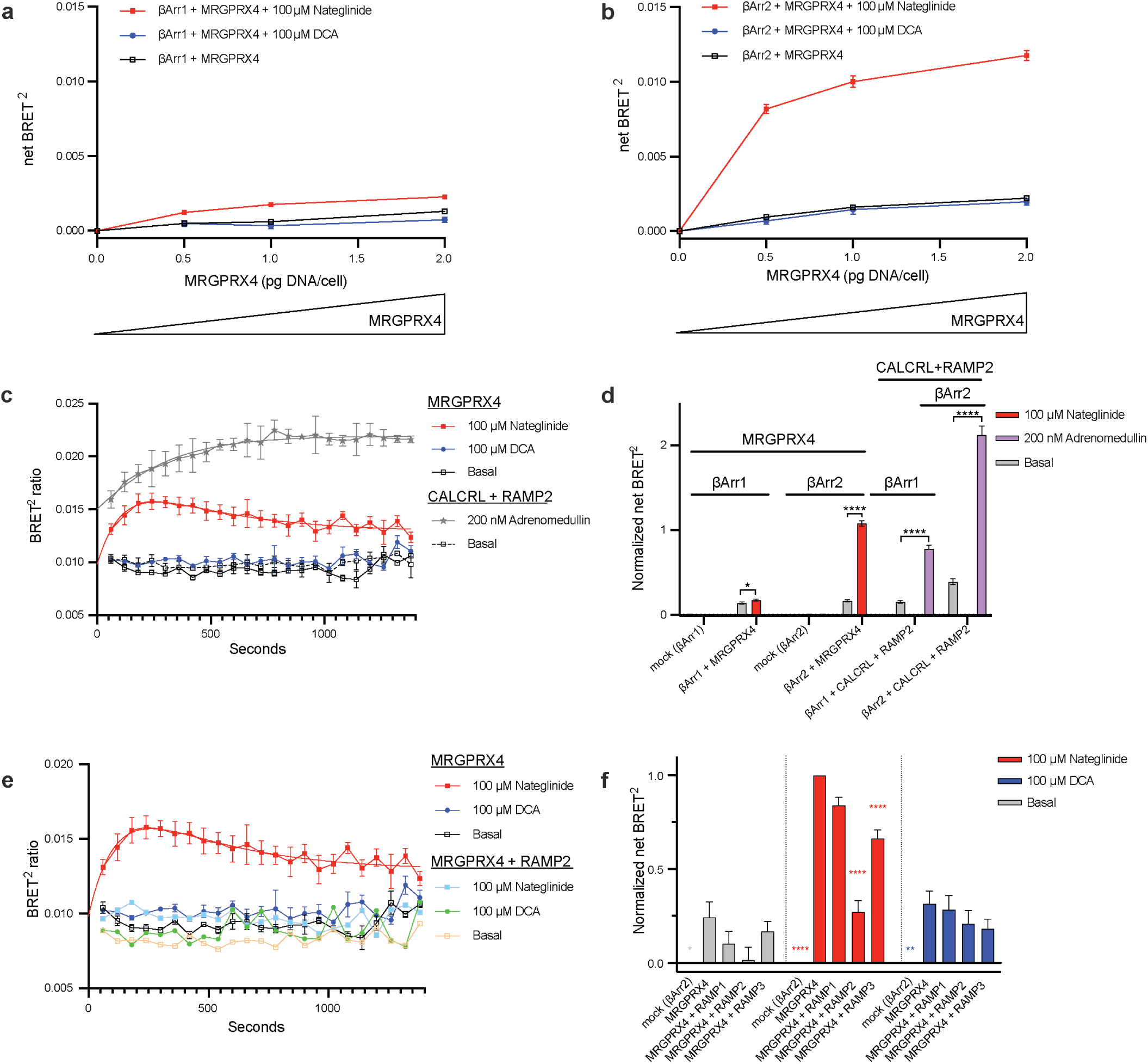
The effect of RAMP co-expression on MRGPRX4 β-arrestin recruitment. **a-b** BRET^2^ β-arrestin recruitment assays were carried out in the presence of increasing amounts of MRGPRX4 co-expressed with β-arrestin1 (**a**) or β-arrestin2 (**b**) upon stimulation with 100 µM nateglinide or DCA. **c** Time course of β-arrestin2 recruitment in cells expressing MRGPRX4 upon stimulation with nateglinide (red) and DCA (blue) as compared to basal levels (open black). Cells co-expressing CALCRL with RAMP2 and stimulated with 200 nM adrenomedullin served as the positive control (gray star). Red and gray smooth curves are fits to a two-phase decay model (see **Supplementary Table 3**). **d** Comparison of normalized net BRET^2^ for β-arrestin1 or β-arrestin2 recruitment to MRGPRX4 upon nateglinide stimulation. Cells co-expressing CALCLR with RAMP2 and stimulated with adrenomedullin served as the positive control. Data are normalized to nateglinide-dependent β-arrestin2 recruitment to MRGPRX4. The statistical significance was determined by unpaired two-tailed t test (see **Supplementary Table 2**). **e** Time course of β-arrestin2 recruitment to MRGPRX4 in cells expressing MRGPRX4 alone or with RAMP2 upon stimulation with nateglinide (red) and DCA (blue). The red smooth curve is the fit to a two-phase decay model (see **Supplementary Table 3**). **f** Comparison of normalized net BRET^2^ for β-arrestin2 recruitment to MRGPRX4 under basal conditions and nateglinide or DCA stimulation. MRGPRX4 was expressed alone or co-expressed with each RAMP. Data are normalized to nateglinide-dependent β-arrestin2 recruitment to MRGPRX4. The statistical significance was determined by ordinary one-way ANOVA followed by Dunnett’s multiple comparisons test to nateglinide-stimulated MRGPRX4 (see **Supplementary Table 2**). (\*\*\*\**p* < 0.0001, \*\**p* < 0.01, \**p* < 0.05, if not marked then not significant). Error bars signify the mean ± SEM. For **a, b, d,** and **f** data are from three independent experiments with three replicates each, except for mock (**d** and **f**), which had two replicates per experiment. For **c, e** data are from three independent experiments with two replicates each, except the CALCRL-RAMP2 dataset in (**c**) for which the data are from two independent experiments with two replicates each.

Finally, we tested the effect of RAMP co-expression on β-arrestin2 recruitment to MRGPRX4 (**Fig. 5e,f**). Comparing the effect of all three RAMPs, RAMP1 co-expression did not have a noticeable effect (∼10%). RAMP2 co-expression resulted in the most striking decrease in β-arrestin2 recruitment upon nateglinide treatment (73%). RAMP3 co-expression resulted in a minor attenuation of recruitment (34%). None of the RAMPs had a significant effect on β-arrestin2 recruitment to MRGPRX4 upon DCA treatment or in basal conditions (**Fig. 5f, Supplementary Table 2**). Further, RAMP2 co-expression resulted in near complete suppression of time-dependent β-arrestin2 recruitment to MRGPRX4 in response to nateglinide, and no change for DCA (**Fig. 5e, Supplementary Table 3**). These data show that MRGPRX4 recruits β-arrestin2 but not β-arrestin1, with a profound ligand bias towards nateglinide. RAMP2 co-expression almost fully abolished nateglinide-dependent β-arrestin2 recruitment.

### RAMP2 co-expression decreases MRGPRX4 surface expression

MRGPRX4-RAMP2 complex formation correlated with an attenuation of basal- and agonist-dependent Gq activation and β-arrestin recruitment. To investigate whether RAMP co-expression affects MRGPRX4 total expression and surface expression, we developed a quantitative nanoBRET pulse-chase surface labeling assay (**Fig. 6a**). The assay relies on Tet-On inducible MRGPRX4 N-terminally tagged with NanoLuc luciferase (NLuc) and Halotag 7 (HT7). First, we characterized the functionality of Tet-On NLuc-HT7-MRGPRX4. Employing the IP1 accumulation assay, we validated that the receptor responds to nateglinide and DCA similarly to all previously used MRGPRX4 constructs (**Supplementary Fig. 4a-f, Supplementary Fig. 5**). Further, the pharmacological effects of the RAMPs on Tet-On MRGPRX4 are in line with previous characterization (**Supplementary Fig. 5d-h, Fig. 4**). We optimized the assay conditions by testing different levels of MRGPRX4 expression and induction by treatment with doxycycline (dox) and reading out IP1 accumulation or total expression by NanoGlo Luminescence (**Supplementary Fig. 5a-c, i**).

**Fig. 6.**
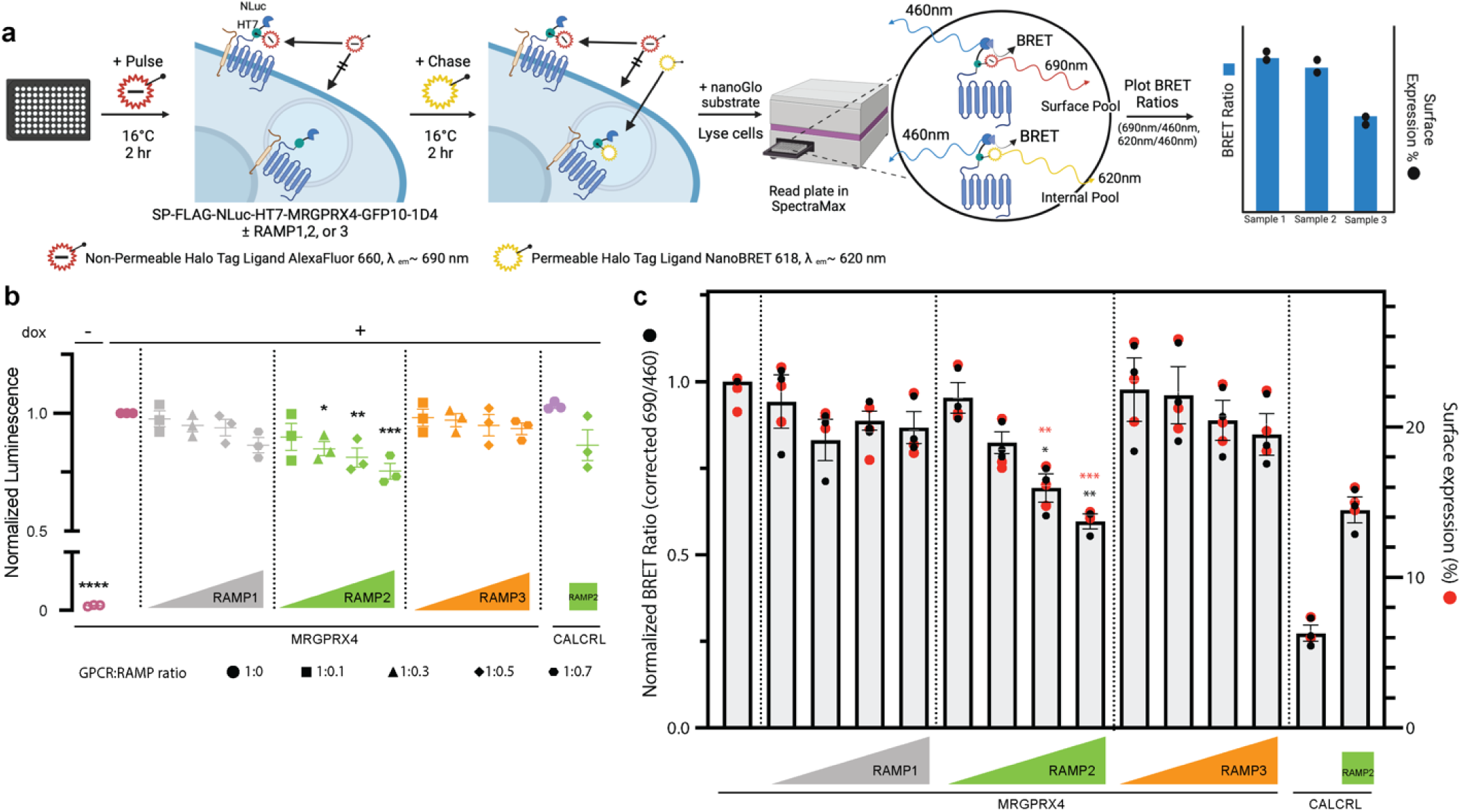
The effect of RAMP on total expression and surface expression of MRGPRX4. **a** Schematic cartoon of NanoBRET pulse-chase experiment design. **b** NanoBRET assay was carried out in cells expressing Tet-On MRGPRX4 alone or co-expressed with increasing amounts of each RAMP. Tet-On CALCRL expressed alone and co-expressed with RAMP2 were included as controls. Total receptor expression is plotted as total luminescence normalized to the MRGPRX4 (+ dox) condition and is from three independent experiments with six replicates each. **c** Surface labeling of MRGPRX4 is plotted as the BRET ratio (corrected 690 nm/460 nm ratio) normalized to MRGPRX4 (left axis, gray bars and black circles). The overlaid red circles show the respective percentage of surface expression for each condition (right axis). Error bars signify mean ± SEM and are from three independent experiments with three replicates each. Statistical significance was determined by ordinary one-way ANOVA followed by Dunnett’s multiple comparisons test to MRGPRX4 expressed alone for total expression of MRGPRX4 (**b**), surface labeling (**c**, black *) and surface expression percentage (**c**, red *) (see **Supplementary Table 2**) (*\*\*\*p* < 0.001, \*\**p* < 0.01, \**p* < 0.05, if not marked then not significant). Schematic created with BioRender.com.

Next, we proceeded with the nanoBRET assay, which employed a cell-impermeable HT7 ligand for the pulse and a cell-permeable HT7 ligand for the chase. The pulse step labeled all surface expressed MRGPRX4, while the chase step labeled any remaining MRGPRX4. The pulse and chase labeling could be measured simultaneously and deconvoluted to determine total and surface expression (**Fig. 6a**). We measured total MRGPRX4 expression with increasing levels of RAMP co-expression (**Fig. 6b, Supplementary Table 2**). As expected, MRGPRX4 expression was dox-dependent. RAMP1 and RAMP3 co-expression did not alter MRGPRX4 total expression at any level. On the other hand, we observed a small reduction in MRGPRX4 total expression, up to 25%, that scaled with the level of RAMP2 (**Fig. 6b, Supplementary Fig. 5i**). CALCRL expressed alone and CALCRL expressed with RAMP2 were the positive controls (**Fig. 6b**).

Finally, we assessed MRGPRX4 surface expression with increasing amounts of RAMP (**Fig. 6c, Supplementary Table 2**). RAMP1 or RAMP3 co-expression did not significantly affect MRGPRX4 surface expression. Interestingly, we observed a strong attenuation of MRGPRX4 surface expression when it was co-expressed with RAMP2, an effect that scaled with the amount of RAMP2 expressed. CALCRL expressed alone or with RAMP2 served as a negative and positive control, respectively, and as expected CALCRL surface expression strongly increased upon RAMP2 co-expression^14, 15^ (**Fig. 6c**). Together, these data show that co-expression of MRGPRX4 with RAMP2, but not RAMP1 or RAMP3, strongly attenuates MRGPRX4 surface expression.

### Prediction of MRGPRX4-RAMP2 complex structure with AlphaFold Multimer

To complement the experimental results, we employed AlphaFold Multimer to predict the MRGPRX4-RAMP2 complex structure *in silico*^16, 17^. The MRGPRX4-RAMP2 structural model shows that the three extracellular α helices of RAMP2 cap the extracellular side of MRGPRX4. The RAMP2 transmembrane (TM) α helix lies close to TM5 and TM6 of MRGPRX4 (**Fig. 7a**). The disulfide between TM4 and TM5 observed in the solved MRGPRX4 structure (and in the solved MRGPRX1 structures) also exists in the AlphaFold prediction^18, 19^. Based on the Predicted Aligned Error (PAE), which is used to assess inter-protein model confidence, the confidence of the prediction of the three extracellular α helices of RAMP2 relative to MRGPRX4 was medium-high, but that of the RAMP2 transmembrane α helix to MRGPRX4 was low (**Fig. 7b**). This may be in part because AlphaFold is not trained with membranes and does not include membranes in the prediction. Nonetheless, the reliability of AlphaFold2, the latest iteration of AlphaFold, even in membrane proteins has been studied and found trustworthy^20^. Next, we used PDBePISA to list the predicted interacting residues between MRGPRX4 and RAMP2 and then checked each pair manually in ChimeraX^21, 22^. We found that the predicted interface area of MRGPRX4 is 971 Å^2^ (**Fig. 7c**). There are three potential salt bridges between MRGPRX4 and RAMP2, in addition to hydrogen bonds to main chain atoms. Most of the van der Waals interactions are contained in the TM region (**Fig. 7d**, **Supplementary Table 4**). Although the ECD of RAMP2 appears to “cap” MRGPRX4, only two predicted RAMP2-interacting residues are also reported to contribute to the MRGPRX4 binding pocket (Arg95 and Lys96, **Supplementary Fig. 6**)^18^.

**Fig. 7.**
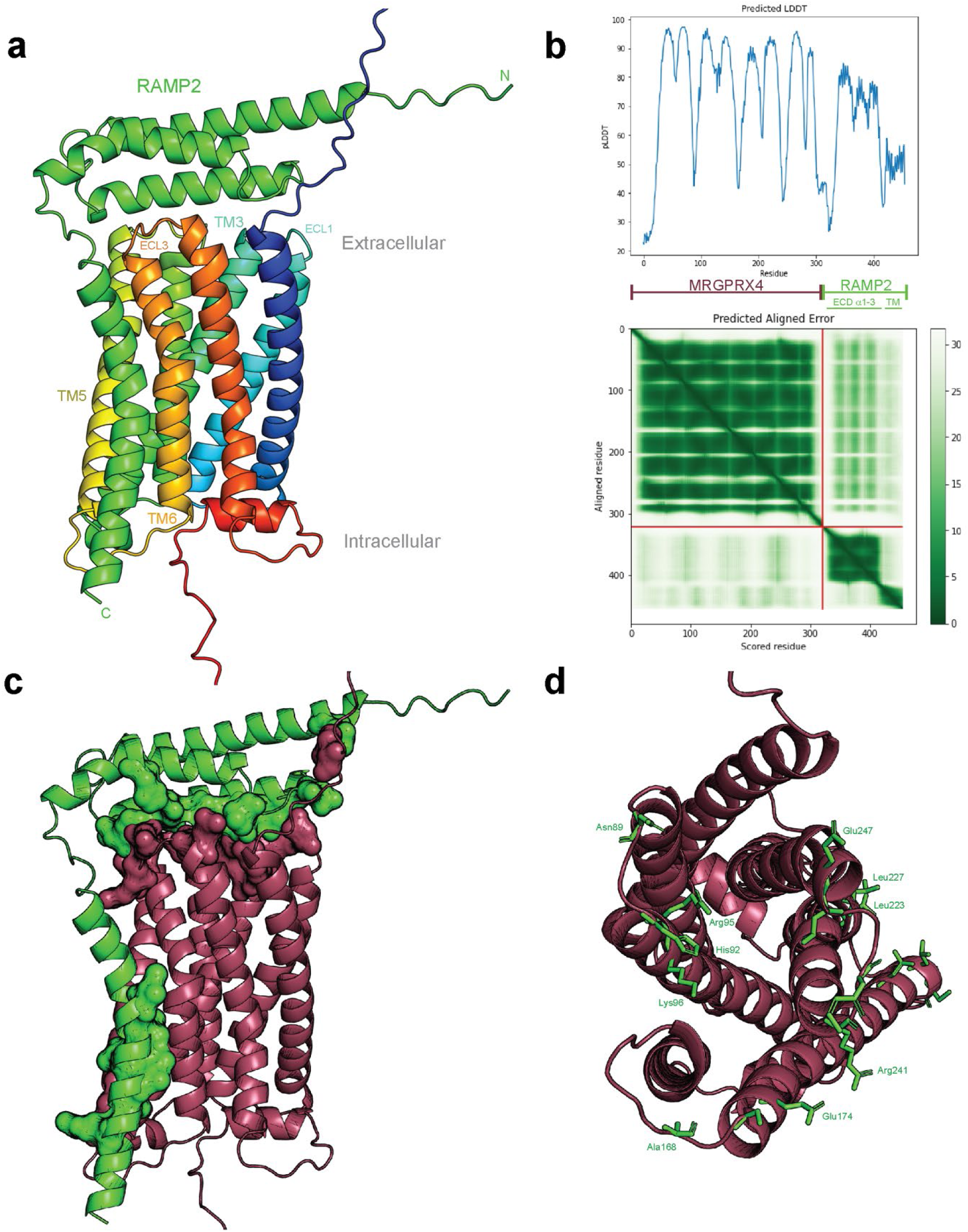
Alpha-Fold Multimer prediction of MRGPRX4-RAMP2 complex structure. **a** Predicted complex formation between MRGPRX4 and RAMP2 generated with Alpha-Fold Multimer Colab^16, 17^. MRGPRX4, rainbow color gradation from deep blue (N-terminal tail) to deep red (C-terminal tail); RAMP2, green. ECL, extracellular loop; TM, transmembrane domain. **b** Predicted local distance difference test (pLDDT) plot (top) and Predicted Aligned Error (PAE) two-dimensional plot (bottom), generated by Alpha-Fold Multimer Colab. **c** Predicted complex structure of MRGPRX4-RAMP2 with the interacting residues shown as surfaces. The predicted interacting residues were calculated by PDBePISA. MRGPRX4, maroon; RAMP2, green. **d** Top view of MRGPRX4 from extracellular side, with predicted RAMP2-interacting residues shown as green sticks. MRGPRX4, maroon. The full list of predicted interacting residues is provided in **Supplementary Table 4**.

## Discussion

MRGPRX4 was recently deorphanized and described as an itch receptor^2–4^, but relatively little is known about its function in different physiological contexts. We hypothesized that RAMPs may be a missing component in our understanding of MRGPRX4 and applied an SBA assay to test for MRGPRX4-RAMP interactions. MRGPRX4 is not the first class A GPCR that has been shown to interact with RAMPs^15^, however it is the first delta subfamily class A GPCR with RAMP interactions that have been both identified and functionally characterized. The gamma subfamily class A GPCR atypical chemokine receptor 3 (ACKR3) was recently shown to interact with the RAMPs, with distinct functional consequences for the ACKR3-RAMP3 interaction^23^. Interestingly, in the case of MRGPRX4, although both RAMP2 and RAMP3 appear to interact with MRGPRX4, we observed marked functional consequences for only MRGPRX4-RAMP2 complexes, as manifested by a strong attenuation of both the basal and agonist-dependent Gq activity for all agonists tested. Based on this striking observation, we hypothesized that RAMP2 may also affect β-arrestin recruitment to MRGPRX4 and therefore impart MRGPRX4 receptor bias. Indeed, if the presence of RAMP2 was correlated with an increase of β-arrestin recruitment to MRGPRX4, it would explain why RAMP2 co-expression suppresses MRGPRX4-mediated Gq activation. However, we found that MRGPRX4-RAMP2 complex formation correlated with decreased β-arrestin recruitment after agonist treatment and a trend towards decreased recruitment in basal conditions. Therefore, the interaction between MRGPRX4 and RAMP2 causes a decrease in Gq activation and β-arrestin recruitment, both of which may be explained by the observation that MRGPRX4 surface expression and total expression are attenuated by RAMP2 (**Fig. 8**).

**Fig. 8.**
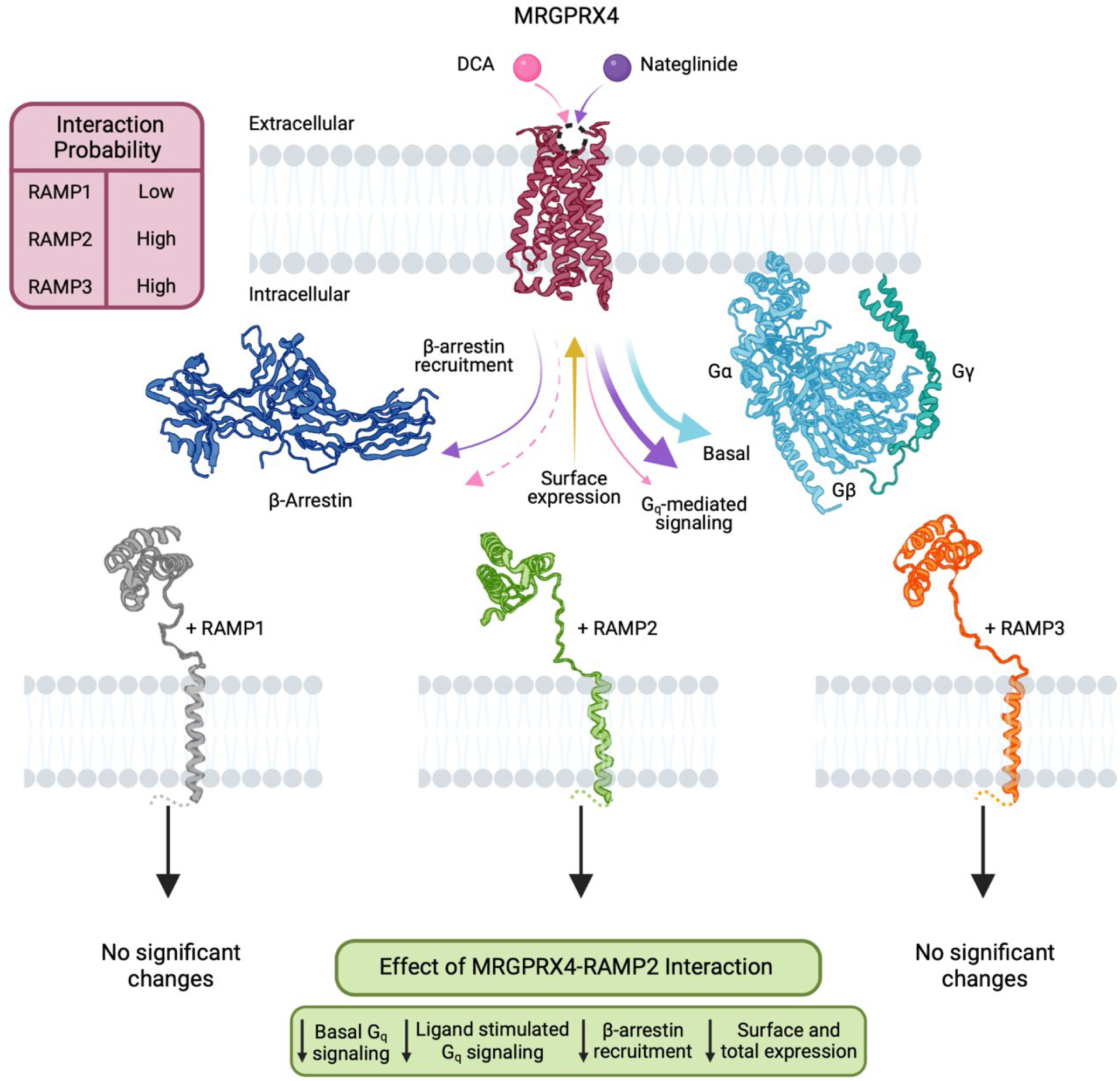
Summary of identified MRGPRX4-RAMP complexes and functional consequences. Schematic summarizing the DCA-stimulated, nateglinide-stimulated, and basal Gq-mediated signaling of MRGPRX4, and the corresponding β-arrestin recruitment. The maroon table indicates interaction probability between MRGPRX4 and each RAMP as determined by the statistical significances calculated from results of the SBA immunoassay and the PLA. The observed effects of RAMP2 co-expression on MRGPRX4 are summarized in the lime green table. Of the three RAMPs, only RAMP2 co-expression with MRGPRX4 correlated with a significant decrease of all parameters measured. Structures and symbols are as follows: DCA (pink ball), nateglinide (purple ball), MRGPRX4 (maroon; PDB 7S8P), Gq protein (α and β, light blue; γ, turquoise; PDB 1GP2), β-arrestin2 (navy blue; PDB 1G4M), RAMP1 (gray; PDB 6E3Y), RAMP2 (lime green; PDB 6UUN), RAMP3 (orange; PDB 6UUS), membrane lipids (light gray). Colored arrows represent the basal (light blue), DCA-induced (pink), and nateglinide-induced (purple) Gq signaling and β-arrestin recruitment promoted by MRGPRX4, and the receptor cell surface expression (ocher). Relative thickness of the arrows corresponds to strength of the elicited responses. Dashed line indicates that the response was measured but not observed. Schematic created with Biorender.com.

RAMP2 has been reported to have pleiotropic effects on G protein activation and β-arrestin recruitment^15^. For example, it promotes β-arrestin recruitment to CALCRL and parathyroid hormone 1 receptor (PTH1R) but has no effect on β-arrestin recruitment to receptors such as the secretin receptor (SCTR)^15, 24–26^. Conversely, RAMP2 co-expression with the glucagon receptor (GCGR) abolishes β-arrestin recruitment, promotes enhanced basal and agonist-dependent GCGR internalization, and attenuates Gq activation while increasing intracellular Gs activation^27, 28^. Although RAMP2 was originally described as a chaperone, specifically for CACLRL, it can also act as an “anti-chaperone” in some instances, such as for GCGR, causing a receptor that would otherwise localize to the cell surface to remain intracellular. We observed a similar result with RAMP2 and MRGPRX4.

Comparing the recently solved cryogenic electron microscopy (cryo-EM) structure of MRGPRX4 to structures of CALCRL-RAMP complexes not surprisingly reveals significant differences between MRGPRX4 and CALCRL, particularly in the extracellular domain, suggesting that the RAMP mode of binding and regulation of MRGPRX4 may be distinct^14, 18, 29, 30^. An open question remains regarding the ratio of free MRGPRX4 to RAMP2-bound MRGPRX4 on the cell membrane and intracellularly, as well as the precise kinetics, dynamics and stoichiometry of complex formation. Further, we do not yet know which intracellular compartment MRGPRX4 localizes to and how the RAMPs affect its organelle-specific localization pattern.

Our finding that RAMP2 causes attenuation of MRGPRX4 total expression levels may represent another mode of MRGPRX4 regulation by RAMP2. RAMP2 may attenuate MRGPRX4 expression by altering its cellular localization, resulting in the cell detecting an improperly localized (intracellular) receptor and marking it for degradation^31^. An alternative and somewhat less probable hypothesis is that RAMP2 may promote the ubiquitination of MRGPRX4, thereby leading to its degradation by the proteosome. MRGPRX4 has three lysine residues within its intracellular tail. However, none of these have been shown to be ubiquitination sites. RAMP2 has one C-terminal lysine, but there are conflicting reports regarding whether it is a ubiquitination site or not^15^.

In contrast to RAMP2, RAMP3 appears to form complexes with MRGPRX4, but does not cause a significant change to Gq signaling, β-arrestin recruitment, total expression, or surface expression. Therefore, the modulatory role of RAMP3, if any, on MRGPRX4 is still unknown. RAMP3 has a unique C-terminal tail motif compared with the other RAMPs and has been shown to affect the downstream trafficking of several interacting receptors, such as CALCRL and the atypical chemokine receptor 3, ACKR3^15, 23, 32–34^. Therefore, the elucidation of the mode of MRGPRX4 desensitization, degradation and recycling in the presence or absence of each RAMP may reveal unique effects of RAMP3.

We did not observe MRGPRX4 agonist signaling bias associated with co-expression of any of the three RAMPs. If the intracellular retention of MRGPRX4 is the primarily driver for the effect of RAMP2 on both basal and agonist-dependent Gq activation, a RAMP2-associated agonist bias is not to be expected. On the other hand, MRGPRX4 appears to exhibit agonist bias for its two most potent activators tested, nateglinide and DCA. Based on the pharmacological characterization of MRGPRX4 agonists, we focused on nateglinide and DCA for subsequent studies for two reasons. First, nateglinide and DCA were the most potent activators of MRGPRX4. Second, here and in previous reports, DCA was shown to activate the same pathway as other BAs, but more potently^2, 4^. By comparing the MRGPRX4 pharmacology of nateglinide to DCA, it may be reasonable to extend the trends observed for DCA to UDCA, TDCA and other BAs.

We show here that nateglinide and DCA activate MRGPRX4 through the same Gq-PLC signaling pathway, albeit to different extents. Both nateglinide and BA-mediated MRGPRX4 Gq signaling have been previously reported. However, the results across reports are not always consistent. Yu and colleagues showed that DCA activates MRGPRX4-promoted Ca^2+^ flux somewhat more potently than nateglinide, although no EC_50_ values were reported. These results, which are not consistent with ours, also do not align with the findings of Cao and colleagues, who determined the Ca^2+^ flux EC_50_ of nateglinide to be 4.717 µM^4, 18^. The original study uncovering MRGPRX4 activation by nateglinide reports a phosphatidylinositol (PI) hydrolysis EC_50_ of 2.1 µM^10^. The EC_50_ discrepancies may reflect differences in the Gq second messenger assay employed or experimental conditions, such as transfection characteristics (stable versus transient MRGPRX4 expression), the tags on the MRGPRX4 construct used, or assay properties. We have extensively characterized the MRGPRX4 constructs we used in this study, and validated functionality for all of them. Here, by employing an assay that measures the accumulation of a different downstream second messenger, IP1, which is a degradation product of the second messenger D-myo-inositol-1,4,5-trisphosphate, we show that nateglinide is a consistently more potent Gq activator than DCA. Moreover, our IP1 accumulation EC_50_ for nateglinide is within an order of magnitude of that calculated by Cao et al. and Kroeze et al. employing different second messenger assays^10, 18^.

We show that both YM and U73122 inhibit the MRGPRX4-mediated IP1 accumulation upon DCA or nateglinide treatment, suggesting that both nateglinide and BAs activate the same Gq-PLC signaling pathway through MRGPRX4 as reported earlier^2, 4^. Nateglinide activates both Gq and β-arrestin recruitment, while DCA is biased towards Gq and away from β-arrestin. Nateglinide treatment of MRGPRX4 activates Gq 48% more potently than DCA. On the other hand, while nateglinide induces β-arrestin2 recruitment to MRGPRX4, DCA does not. The difference between the two agonists is much more pronounced in terms of β-arrestin2 recruitment than Gq signaling. Nateglinide activates β-arrestin2 recruitment to MRGPRX4, but the recruitment is lower compared to CALCRL-RAMP2. This observation is not surprising, considering that MRGPRX4 belongs to the class A, or rhodopsin family, whereas CALCRL is a class B (secretin family) GPCR that is known to stably recruit β-arrestins^35^. A thorough characterization of the CALCRL-RAMP β-arrestin recruitment profiles was recently published and showed that CALCRL in complex with all three RAMPs can competently recruit both β-arrestins^36^. Although not measured with our BRET assay, MRGPRX4 may indeed interact with β-arrestin2 upon DCA stimulation but exist in a much more transient ensemble of conformations than after nateglinide stimulation. Therefore, desensitization of MRGPRX4 upon DCA stimulation may occur through an alternative pathway, for example through an arrestin domain-containing protein (ARRDC)^37^, or through MRGPRX4 interacting with a specific GPCR kinase (GRK), as was shown to be the case for the leukotriene B4 receptor, BLT1^38, 39^. To the best of our knowledge, this is the first characterization of the β-arrestin recruitment to MRGPRX4 outside of the β-arrestin recruitment PRESTO-Tango assay, which does not reflect the ability of MRGPRX4 to recruit β-arrestins^10^. Moreover, we report here for the first time a signaling bias of a bile acid at MRGPRX4.

The discovery of novel MRGPRX4-RAMP2 and MRGPRX4-RAMP3 complex formation is physiologically relevant as human skin keratinocytes, neurons in the TG, and neurons in the DRG are known to express both MRGPRX4 and the RAMPs^5–9, 40–44^. MRGPRX4 is primarily expressed in small-diameter sensory neurons of the DRG and TG and is also expressed in human skin keratinocytes^6–9^. Interestingly, skin keratinocytes were found to express RAMP1 and RAMP2, but not RAMP3^40^. In contrast, all three RAMPs are expressed in the TG and in the DRG^41–44^. RAMP2 and RAMP3 seem equally competent to form complexes with MRGPRX4, however based on what we have shown, RAMP2 is the only biologically relevant MRGPRX4 interactor. The co-expression of MRGPRX4 and RAMP2 in the DRG is likely key for mediating cholestatic itch but their co-expression in skin keratinocytes may also contribute to the itch propagation mechanism. Whether RAMP3 may prevent RAMP2 from forming complexes with MRGPRX4 in certain physiological or pathophysiological contexts such as in the TG and DRG but not in skin keratinocytes is unknown. More broadly, whether the RAMPs compete with each other is an open question for the field as a whole. Moreover, MRGPRX4 is co-expressed with histamine receptor 1 (HRH1) in DRG neurons, but potential heterodimerization of the two receptors has not been studied^4, 45^. The RAMPs may further regulate MRGPRX4 by promoting or disrupting its heterodimerization.

As MRGPRX4 is expressed in several cell types, the consequence of RAMP interaction may vary accordingly. Therefore, studying MRGPRX4 signaling mediated by BAs in different contexts will shed more light on whether BAs are indeed the pruritogens for cholestatic itch, a conclusion which is not yet fully accepted by all in the field^46^. It will be interesting to see if further studies of cholestatic itch in humans reveal physiological differences that correspond to the differences in pharmacology at the MRGPRX4 receptor. Moreover, measuring the expression of both MRGPRX4 and RAMP2 in the relevant tissues of subjects may help explain variability in the itch response data, as plasma bile levels do not correlate well with presentation of itch^1, 47, 48^. MRGPRX4 also displays a high level of single nucleotide polymorphisms (SNPs)^1, 49, 50^. For example, two MRGPRX4 variants expressed exclusively within African American participants are associated with a preference for menthol cigarettes and correlated with reduced signaling in cell-based assays^51^. Missense SNPs, particularly the four SNPs with an allele frequency of >20%, may affect the ability of MRGPRX4 to form complexes with RAMP2^1^.

Other GPCRs are also shown to modulate itch response, including the Mas-related family members MRGPRX1 and MRGPRX2^1, 50, 52–54^. The four MRGPRX subtypes (X1-X4) are closely related and, except for orphan MRGPRX3 for which very little is known, they all are considered itch receptors. MRGPRX4 is most closely related to MRGPRX3 and is most distant from MRGPRX2 within the “X” branch of the MRGPR phylogeny^1^. The discovery of MRGPRX4-RAMP interactions raises the intriguing possibility that other itch receptors from the same family may also interact with the RAMPs. If phylogenetic similarity is an indicator for probability for RAMP interaction, MRGPRX3 and MRGPRX1 are the most likely candidates. It will be interesting to interrogate whether the MRGPRX structural differences result in altered RAMP-binding ability of the receptors.

Structurally, the MRGPRX family is “non-canonical” in several aspects and does not harbor typical class A GPCR features such as the disulfide bond between TM3 and ECL2. We show here that residues located on ECL1-3 and the extracellular side of TM3 and TM5 of MRGPRX4 constitute the predicted interaction interface with RAMP2 (**Fig.7, supplementary Fig. 6, Supplementary Table 4a**). Compared with MRGPRX4, the structure of the extracellular surfaces of MRGPRX1 and MRGPRX2 are distinct, but all three share the common feature of a shallow binding pocket^18, 19^.

The structure of MRGPRX4 was recently solved in complex with a compound derived from nateglinide, MS47134, that binds through both polar and non-polar interactions^18^. MRGPRX4 has an overall positive electrostatic potential surface on its binding interface and therefore binds anionic agonists like BAs. The observed ligand binding was very close to the extracellular surface, which may explain why MRGPRX4 binds a range of BAs and other agonists^55^. A putative positive allosteric modulator (PAM) binding site on MRGPRX4 was previously suggested for bilirubin^4, 18^. The recently solved structure of MRGPRX1 revealed the presence of distinct putative orthosteric and allosteric pockets^19^, but the structure of MRGPRX4 solved with nateglinide analogue MS47134 did not reveal analogous pockets. Instead, the shallow positively-charged orthosteric pocket was distant from the canonical orthosteric binding site for most class A GPCRs^18^. Taken together, it is likely that both nateglinide and bile acids bind in this shallow anomalous orthosteric site on MRGPRX4. However, further confirmatory studies are needed to define the structural pharmacology and activation mechanism of MRGPRX4.

The positioning of RAMP2 relative to MRGPRX4 in the predicted complex model generated using AlphaFold Multimer seems plausible based on previously solved GPCR-RAMP structures^29, 30, 56, 57^. The positioning of RAMP2 does not clash with G-protein binding, as observed in the cryo-EM structure of Gq-coupled MRGPRX4^18^. Overall, the binding site of the nateglinide analogue used for the structural determination of MRGPRX4 (PDB 7S8P) was distinct from the predicted RAMP2-binding interface. Minimal contact between RAMP2 and the binding pocket is consistent with our experimental data suggesting that RAMP2 is not directly modulating ligand binding. Previous structural studies also provide evidence for RAMPs exerting their modulatory function allosterically^29, 30, 56, 57^.

In summary, we show that MRGPRX4 interacts with RAMPs, and that MRGPRX4 complex formation with RAMP2 specifically alters receptor cell surface expression and total expression. We also show that BAs are biased agonists at MRGPRX4 toward Gq signaling and away from β-arrestin. We employed AlphaFold Multimer to predict a putative complex structure between RAMP2 and MRGPRX4, a delta subfamily class A GPCR. The complex model can be tested in future work using mutagenesis, targeted photo-crosslinking^58–60^, and structural studies. Collectively, these data illustrate a critical role of RAMPs in MRGPRX4 pharmacology and drug development aimed at cholestatic itch, and potentially suggest a more general role of RAMPs in regulating the biology of class A GPCRs.

## Materials and methods

### Reagents

Nateglinide (23320), UDCA (15121), and TDCA (15935) were from Cayman Chemical. DCA (30960), Adrenomedullin (A2327) was from Sigma-Aldrich. CGRP (4013281) was from Bachem. YM254890 (CAS:568580-02-9) was from Wako Pure Chemical Industries. U73122 (70740) was from Cayman Chemical. BRET substrate Prolume Purple (Methoxy e-TZ) (369) was from NanoLight Technology. Halo Tag NanoBRET 618 ligand (G9801), Halo Tag Alexa Fluor 660 Fluorescent ligand (G8471), and NanoGlo luciferase (N1110) were from Promega. Doxycycline was from Clontech (631311). The IP-One HTRF kit was from CisBio (62IPAPEB). Bovine serum albumin (BSA) fraction V, fatty acid-free, was from Roche (9048-46-8). Poly-D-lysine and LiCl were from Sigma-Aldrich. HEK293T cells were from the American Type Culture Collection. HEK293 Freestyle cells were from Thermo Fischer Scientific. Dulbecco’s modified Eagle’s medium (DMEM) GlutaMAX (10564-011), FluoroBrite DMEM (A18967-01), Dulbecco’s PBS (DPBS) (14190144), HEPES buffer (25-060-CI), L-glutamine (25030081), Lipofectamine 2000 (11668019), and Tet-system approved FBS (A4736401) were from Thermo Fisher Scientific. Fetal bovine serum (FBS) (10437528) was from Gemini Bio-Products. *n*-Dodecyl-b-d-maltoside (DM) detergent (D310S) was from Anatrace. DC Protein Assay Kit (5000112) and Precision Plus protein dual standards (1610374) were from Bio-Rad. cOmplete Mini, EDTA free protease inhibitor cocktail (11836170001) was from Sigma-Aldrich. LoBase clear bottom/black small-volume 384-well microplates (788890), microplate lids ultra-low profile (691161), and clear bottom/ black 96-well microplates (polystyrene wells, flat bottom; 655986) were from Greiner Bio-One. NuPage 4-12% Bis-Tris Gel Invitrogen (NP0336BOX), NuPage MES SDS Running Buffer (20x) (NP0002), and Invitrogen NuPage LDS Sample Buffer (NP0007) were from Thermo Fisher Scientific. Immobilon-FL PVDF Membranes (IPFL00010), aprotinin saline solution (A6279), and PMSF (93482) were from Sigma-Aldrich. Blots were imaged and analyzed on a Licor Odyssey M. NEBuilder HiFi DNA Assembly Master Mix (E2621), Dpn1 (R0176S), T4 DNA Ligase (M0202), Q5 Hot Start High-Fidelity DNA Polymerase (M0491S), and dNTPs (N0447S) were from New England BioLabs. Oligonucleotides which are listed in **Supplementary Table 5** were purchased at the standard desalting grade from Integrated DNA Technologies. TagMaster Site Directed Mutagenesis kit was from GM Biosciences. QIAGEN Plasmid Maxi Kits and QIAprep Spin Miniprep Kit were from QIAGEN. Zymo PUREII Plasmid Midiprep kit (D4200) was from Zymo Research. Zymo DNA clean and concentrator kit (D4004) was from Genesee Scientific. NaveniFlex MR (NF.MR.100) PLA kit was from Navinci. Duolink in Situ Mounting Medium with DAPI (DUO82040) was from Sigma-Aldrich. Rabbit anti-β-actin primary antibody was from Thermo Fisher (PA1-16889). Rat anti-OLLAS and mouse anti-1D4 antibodies were in house. PE conjugation kit was from Abcam (102918). Rabbit anti-HA antibody (C29F4) was from Cell Signaling Technology. Monoclonal anti-FLAG M2 antibody (F3165) was from Sigma-Aldrich. PE conjugated anti-rabbit IgG and PE conjugated anti-mouse IgG were from Jackson ImmunoResearch. Secondary antibodies Goat anti-rat 800CW, Goat anti-rabbit 680RD (926-68071), and Goat anti-mouse 800CW (926-32210) were from LI-COR Biosciences. PE-conjugated rat anti-FLAG (637309), PE-conjugated mouse anti-HA (901517), and mouse anti-HA (16B12) were from BioLegend. Sheep anti-RAMP1, -RAMP2, and - RAMP3 (AF6428, AF6427, and AF4875, respectively) were from R&D Systems. Mouse IgG antibody (PMP01X) was from Bio-Rad. Rabbit IgG antibody (P120-101) was from Bethyl Laboratories. Dithiothreitol (CAS:27565-41-9) was from Gold Biotechnology. ProClin 300 (48912-U), casein (C7078), polyvinyl alcohol (PVA) (25213-24-5), Polyvinylpyrrolidone (PVP) (9003-39-8) were from Sigma-Aldrich. Blocking Reagent for ELISA (BRE) (11112589001) was manufactured by Roche. 1-ethyl-3-(3-dimethylaminopropyl)-carbodiimide hydrochloride (EDC) (c1100) was from ProteoChem. N-hydroxysuccinimide was from Pierce (CAS:6066-82-6).

### Molecular biology

The primers used for all the molecular biology were designed using the NEBuilder Assembly Tool on the NEB website, purchased from Integrated DNA Technologies, and are listed in **Supplementary Table 5**. The HiFi assembly procedure was performed as previously described^61^. HiFi DNA Assembly was used to generate the Tet-On Inducible Gene Expression vector plasmids SP-FLAG-NLuc-HT7-MRGPRX4-GFP10-1D4 and SP-FLAG-NLuc-HT7-CALCRL-GFP10-1D4 from three parts: a Tet-On Inducible Gene Expression backbone from the construct NLuc-HT7-CysLTR2-GFP10-1D4 generated within the lab; signal peptide (SP), N-terminal FLAG tag, Nano Luciferase (NLuc), and halo tag 7 (HT7) from SP-FLAG-NLuc-HT7-CCR5-CLIP-2xOLLAS-1D4, which is based on a previously published set of constructs^62^; and the GPCR construct of interest C-terminally tagged with GFP10 and 1D4. The original Tet-On vector was designed and purchased from Vectorbuilder. All constructs were confirmed by sequencing in the forward and reverse directions (T7, BGHR primers) (Genewiz).

#### Constructs

***MRGRPX4***: The different complementary DNAs (cDNA) constructs encoding the MRGPRX4 receptor were generated as follows. The mammalian expression pcDNA3.1(+) vector encoding for epitope-tagged human MRGPRX4 cDNA includes an engineered N-terminal HA tag (YPYDVPDYA) and a C-terminal 1D4 tag (TETSQVAPA) (HA-MRGPRX4-1D4). The MRGPRX4 cDNA sequence was codon optimized for expression in human cell lines (Genewiz). The FLAG-MRGPRX4-GFP10-1D4 cDNA construct was generated by inserting the SP-FLAG-MRGPRX4 cDNA sequence from the PRESTO-tango library^10^, where an HA SP sequence (MKTIIALSYIFCLVFA) is followed by a FLAG tag (DYKDDDD) and the receptor sequence, into a GFP10-1D4 containing pcDNA3.1(+) vector obtained from a CysLTR2-GFP10-1D4 constrct^61^. The MRGPRX4-GFP10-1D4 was then inserted into a Tet-On Inducible Gene Expression vector to generate SP-FLAG-NLuc-HT7-MRGPRX4-GFP10-1D4. ***CALCRL***: The codon-optimized sequence of epitope-tagged human HA-CALCRL-1D4 was encoded in pcDNA3.1(+) vector (full 1D4 sequence, DEASTTVSKTETSQVAPA)^13^. The 23 amino acid SP sequence (MRLCIPQVLLALFLSMLTGPGEG) from 5-hydroxytryptamine receptor 3a receptor (5-HT3a) was added to the CALCRL cDNA in place of the native signal sequence, which was determined using SignalP 4.1, as previously described^13^. Then, a GFP10 was inserted into HA-CALCRL-1D4 to generate HA-CALCRL-GFP10-1D4. The CALCRL-GFP10-1D4 was then inserted into a Tet-On Inducible Gene Expression vector to generate FLAG-NLuc-HT7-CALCRL-GFP10-1D4. ***RAMPs*:** Epitope-tagged human RAMP cDNA constructs were encoded in pcDNA3.1(+) expression vector. The human RAMP1, RAMP2, and RAMP3 cDNAs encoded either a N-terminal FLAG tag (DYKDDDDK) or 3xHA tag (YPYDVPDYA) following the signal sequence (amino acids 1 to 26, 1 to 42, and 1 to 27 for RAMP1, RAMP2, and RAMP3 respectively) and two C-terminal OLLAS tags (SGFANELGPRLMGK) separated by a linker (WSHPQFEKGGGSGGGSGGGSWSHPQFEK). The RAMP cDNA sequences were codon optimized for expression in human cell lines. The FLAG-RAMP-OLLAS constructs have been characterized previously^13^. 3xHA-RAMP-OLLAS was generated based on the FLAG-RAMP-OLLAS constructs using the TagMaster site-directed mutagenesis kit according to the manufacturer’s instructions for “long-range mutation”. ***β-arrestin1 and 2*:** β-arrestin1-RLuc3 and β-arrestin2-RLuc3 have been previously described^63^.

### Cell culture and transfection

#### Culture of Human embryonic kidney (HEK) 293T cells

HEK293T cells were cultured in DMEM GlutaMAX supplemented with 10% FBS at 37°C with 5% CO_2_. Cells were transiently transfected directly ‘in plate’ with 1 pg of MRGPRX4 cDNA per cell, unless otherwise specified. Total DNA amount was maintained constant with empty vector pcDNA3.1(+). All transfection reagent mixtures were performed in FluoroBrite DMEM (Live Cell Fluorescence Imaging Medium, without phenol red), and transfected cells were maintained in supplemented FluoroBrite DMEM [15mM HEPES, 4mM glutamine and 10% FBS, or 10% TET-approved FBS for transfection with a Tet-On plasmid]. Briefly, the appropriate amount of plasmid DNA was diluted with FluoroBrite DMEM. In a separate mixture, a volume of Lipofectamine 2000 proportional to 2.5 µL Lipofectamine 2000 per µg of DNA was diluted in FluoroBrite DMEM and incubated for 5 minutes prior to being mixed with the DNA mixture and incubated for 20 minutes. Concurrently, cells were trypsinized, resuspended in 2x supplemented FluoroBrite DMEM, and counted. Cells were mixed with the DNA-Lipofectamine 2000-FluoroBrite DMEM mixture and directly plated onto a black, clear-bottom, tissue culture treated microplate at the cell density of 5,600 cells in 7 µL/well in low volume, LoBase 384-well plates (IP1 accumulation assays), 40,000 cells in 100 µL/well in 96-well plates (BRET^2^ assays), 75,000 cells in 50 µL/well in 96-well plates (NanoBRET assays with Tet-On plasmid), 500,000 cells in 1 mL in 6-well plates (PLA assays) and 1,000,000 cells in 1 mL in 6-well plates (Immunoblot). All microtiter plates were ozone treated and 0.01% poly-D-lysine–coated.

#### Culture of HEK 293 Freestyle (293F) cells

HEK293F cells were cultured in serum-free FreeStyle 293 Expression media using 125 mL disposable culture flasks (Thermo Fisher Scientific). Cells were shaken constantly at 125 rpm at 37°C with 5% CO_2_. Transient transfections were performed using FreeStyle MAX Reagent (Thermo Fisher Scientific) according to the manufacturer’s instructions, and as described previously^13^. The day prior to transfection, cells were diluted to 600,000 cells/mL. The next day, 3 mL of cells were added per well of a 6-well plate. Each well of cells was transfected with 0.5 µg of the indicated RAMP plasmid DNA and/or 0.5 µg of MRGPRX4 plasmid with 3 µL of FreeStyle MAX Reagent. Total transfected plasmid DNA was kept constant at 3 µg with empty vector pcDNA3.1(+).

### Cell lysate preparation

HEK293F cells (for SBA clarified lysate preparation) and HEK293T cells (for expression analysis by Western Blot) were solubilized with *n*-Dodecyl-β-d-maltoside (DM) detergent (Anatrace) to form micelles around membrane proteins and maintain GPCR and RAMP structure and complex formation, as previous described^13^.

### Suspension bead array (SBA) immunoassay

SBA assay was performed to detect MRGPRX4-RAMP complexes from detergent-solubilized lysates and was conducted as previously described^13^. Briefly, antibodies (Abs) were covalently coupled to MagPlex Beads (Luminex Corp). Each Ab was coupled to a unique bead identity^64^. For the SBA assay, clarified HEK293F cells lysates were incubated with an aliquot of the SBA, and protein association with each bead was detected with a PE-conjugated Ab. The fluorescence associated with each bead was measured in a FlexMap 3D instrument (Luminex Corp). The final dilutions used for the detection Abs were 1:1,000 for PE-conjugated anti-FLAG (BioLegend) and PE-conjugated anti-1D4, 1:500 for PE-conjugated anti-OLLAS, and 1:200 for PE-conjugated anti-HA (BioLegend). Two technical replicates with three biological replicates per transfection condition were performed.

For SBA data analysis, data from wells in which there were fewer than 25 beads per ID were excluded. The raw output of median fluorescence intensity (MFI) was then converted to a Robust Z-Score (R.Z-score) across all samples for each capture-detection scheme. An ordinary one-way analysis of variance (ANOVA) with Dunnett’s multiple comparisons test or, for assessing GPCR expression, an unpaired two-tailed t-test, was used to calculate statistical significance compared to mock (Prism 9, GraphPad). The F values and degrees of freedom (DoF) for the ANOVAs, and the t-values and DoF for the t-tests are provided in **Supplementary Table 2**.

### Proximity ligation assay (PLA)

HEK293T cells were transfected as above onto gelatin-coated coverslips within a 6-well plate. After 24 hours cells were fixed as previously described^13^ and then processed following the manufacturer’s instructions for NaveniFlex MR (Navinci) using rabbit anti-HA (Cell Signaling Technology) and mouse anti-FLAG (Sigma-Aldrich) primary Abs at a 1:1500 dilution for each. After PLA processing, cells were mounted in Sigma-Aldrich DuoLink *in situ* mounting medium with 4′,6-diamidino-2-phenylindole (DAPI) (Sigma-Aldrich), allowed to incubate at room temperature (RT) in the dark and imaged the following day.

For PLA image acquisition, deconvoluted PLA images were acquired with a DeltaVision Image Restoration Inverted Olympus IX-71 microscope using a 60x oil immersion objective. Excitation/emission wavelengths are 390 ± 18/435 ± 48 nm for the blue channel (DAPI), 575 ± 25/632 ± 60 nm for the red channel (PLA puncta), and 475 ± 28/ 525 ± 48 for the GFP10 channel (FLAG-MRGPRX4-GFP10-1D4). Exposure times and transmittance percentages were held constant while imaging all samples within the same experiment. Each *Z*-stack image slice is 0.2-µm thick, and each Z-stack was of a different field of view.

Image processing was done in ImageJ (adding scale bars and generating maximum projections) and Imaris. Nuclei stained with DAPI were counted to obtain the total number of cells per image. The PLA puncta were counted in a three-dimensional rendering of each Z-stack in Imaris using the Spot tool. The same Spot parameters (estimated puncta XY and Z diameter, threshold) were used for all samples in all experiments. The puncta count value for each Z-stack was divided by the total number of cells per image, and results were plotted in Prism 9 (GraphPad). Statistics were determined by a one-way ANOVA followed by Dunnett’s multiple comparisons test (Prism 9). The F values and DoF are provided in **Supplementary Table 2**. Outliers from each GPCR-RAMP pair were determined in Prism 9 *via* the ROUT method with *Q* = 1%. Two outliers were removed from the MRGPRX4 dataset and one from the MRGPRX4-RAMP1 dataset.

### IP1 accumulation signaling assays

HEK293T cells were transfected with 1 pg/cell of MRGPRX4 DNA, and the amount of DNA was kept constant at 2 pg/cell with pcDNA3.1(+) empty vector, unless otherwise specified. 24 hours after transfection, IP1 assay was performed as previously described^61^. When applicable, cells were treated with 1 µM YM254890 (YM) or different concentrations of U73122 for 1 hour prior to addition of agonist or buffer. After incubation for 2 hours, Homogenous Time-resolved Fluorescence (HTRF) reagents and IP1 calibration standards were added and incubated for 2 hours in the dark at RT. Time-resolved fluorescence signals were read on the BioTek Synergy NEO-TRF Hybrid multi-mode reader (BioTek Instruments) All data were carried out in three independent experiments with three technical replicate each.

For the agonist dose-response performed with a MRGPRX4 DNA titration, HEK293T cells were transiently transfected with a serial dilution of 2, 1, 0.5, 0.25, 0.125 or 0 pg/cell of FLAG-MRGPRX4-GFP10-1D4 with the total amount of DNA kept constant at 2 pg/cell.

For RAMP DNA titration assay, HEK293T cells were co-transfected with a constant amount of 1 pg/cell of FLAG-MRGPRX4-GFP10-1D4 to which a serial dilution of 1, 0.75, 0.5, 0.25, or 0 pg/cell of each HA-RAMP-OLLAS was added. For a homogenous transfection of MRGPRX4, a master mix of MRGPRX4 DNA was made prior to addition of the RAMP DNA.

Validation of CALCRL constructs was assayed by co-transfecting HEK293T cells with different tagged versions of CALCRL alone or with each RAMP, and with the promiscuous Gqs5 Gq chimera protein, at a respective DNA ratio of 1:1:0.5. Gqs5 is an engineered Gq protein containing the last five amino acid residues of Gs, which allows Gs-coupled GPCRs to signal through Gq downstream signaling pathways^65^. The homogenous transfection method described above was used for all DNA mixes. CALCRL:RAMP2:Gqs5-transfected cells served as positive control for RAMP functionality.

For characterization of the Tet-On Nluc-HT7-MRGPRX4-GFP10-1D4 construct, HEK293T cells were transfected as previously described with some modifications. Doxycycline (dox) dose-response of NLuc-HT7-MRGPRX4-GFP10-1D4 DNA titration was assayed by transfecting HEK293T cells with either 1 or 2pg/cell of NLuc-HT7-MRGPRX4-GFP10-1D4 construct. Receptor expression was induced 4 hours after transfection with addition of different concentrations of dox for 20 hours. For the RAMP DNA titration assay, cells were co-transfected with a constant amount of 1pg/cell of NLuc-HT7-MRGPRX4-GFP10-1D4 DNA and 1, 0.75, 0.5, or 0 pg/cell of each HA-RAMP-OLLAS with the homogenous transfection method. 4 hours after transfection, receptor expression was induced with addition of 1,000 ng/mL dox. 20 hours after induction, IP1 assay was performed.

Data reduction, standard calibration, and transformation of HTRF data were performed as previously described^61^. Normalized IP1 values were calculated relative to the unstimulated mock-transfected cells (set to 0%) and fully stimulated MRGPRX4 (alone if applicable) (set to 100%). These data were fitted to a three parameters sigmoidal dose-response function (Prism 9). The basal and E_max_ parameters describe the lower and upper asymptotic values, respectively, and are listed in **Supplementary Table 1** together with the logEC50, EC50, span and Degrees of Freedom (DoF) values.

### Bioluminescence Resonance Energy Transfer (BRET^2^) β-arrestin recruitment assay

HEK293T cells were transiently co-transfected with β-arrestin-RLuc3 (0.125 pg/cell) and 1pg/cell of FLAG-MRGPRX4-GFP10-1D4 alone or with 1pg/cell of HA-RAMP-OLLAS unless otherwise noted. The method for homogenous transfection of β-Arrestin and BRET^2^ assay of β-Arrestin recruitment has been described previously^61^. Cells were stimulated with BRET buffer with or without 100 µM nateglinide, 100 µM DCA, or 200 nM adrenomedullin. Agonist was incubated for 10 minutes (adrenomedullin) or 3 minutes (nateglinide and DCA) prior to addition of 5 µM of the cell-permeable substrate methoxy e-Coelenterazine (Me-O-e-CTZ/Prolume Purple). BRET^2^ measurements were taken on the BioTek Synergy NEO2 microplate reader.

For MRGPRX4 DNA titration assays, HEK293T cells were co-transfected with either β-Arrestin1 or β-Arrestin2 with increasing amounts of MRGPRX4 (0.5, 1, or 2pg/cell).

To measure the time-course of β-Arrestin2 recruitment to MRGPRX4 and CALCRL, the transfection and general procedure described above was followed, with the modification that Prolume Purple was added first, followed by addition of the appropriate agonists^61^.

The BRET ratio for each sample was determined by calculating the ratio of the light intensity emitted by the GFP10 (515 nm) (acceptor) over the light intensity emitted by the RLuc3 (395 nm) (donor). Net BRET^2^ was determined by subtracting the basal BRET^2^ (β-arrestin-RLuc3 only) signal from the BRET^2^ signals. All BRET values were normalized to the MRGPRX4 only condition. The two-phase decay model fitted parameters for data are summarized in **Supplementary Table 3**. The F values and DoF values are provided in **Supplementary Table 2**.

### Surface labeling assay

HEK293T cells were transfected with 1pg/cell of Tet-On FLAG-NLuc-HT7-MRGPRX4-GFP10-1D4 alone or co-expressed with 0.7, 0.5, 0.3, or 0.1 pg/cell of each RAMP and expression was induced 4 hours after transfection with 1,000 ng/mL dox. HEK293T cells transfected with 1 pg/cell of Tet-On CALCRL alone or with 0.5 pg/cell of RAMP2 served as controls. 20 hours after transfection, media was replaced with BRET buffer. First, the HT7-tagged GPCR was labeled with “pulse” of 100 nM HaloTag Alexa Fluor 660 (cell impermeable, emission 690 nm) for 2 hours at 16°C in the dark. Next, a chase step was performed by adding 100 nM HaloTag 618 (cell permeable, emission 620 nm) for 2 hours at 16°C in the dark. Lastly, Nano-Glo Luciferase assay reagent was prepared per manufacturer’s instructions. Optimal dosage of dox was determined by selecting the concentration at the plateau from a dox dose-response experiment, in which dox-induced expression was assayed with the additional of Nano-Glo Luciferase assay reagent (**Supplementary Fig. 5i**).

Data were acquired by measuring the luminescence at 460, 618, and 690 nm in kinetics mode on a SpectraMax i3X at 37 °C. The obtained emission intensity values were corrected, normalized, plotted against each other, and fitted to a linear equation to generate a scaling factor that was then used to calculate and graph the percentage surface labeling. Experiments were conducted in biological triplicate with three technical replicates each. The corrected 690/460 BRET ratio was normalized to the mean of MRGPRX4 alone (+dox). Similarly, total expression was normalized to the mean of the 460 nm luminescence for MRGPRX4 alone (+dox). The F values and DoF values are provided in **Supplementary Table 2**.

### Immunoblot analysis

Immunoblotting was performed after lysate preparation according to standard procedure. The following abs were used: 1:2,000 Rat anti OLLAS (in house), 1:5,000 Rabbit anti β-actin (Thermo Fisher), 1:4,000 Mouse anti 1D4 (in house), and 1:10,000 (Goat anti Rat 800CW, Goat anti Rabbit 680RD, Goat anti Mouse 800CW; LI-COR). Blots were imaged and analyzed on a LI-COR Odyssey M.

### AlphaFold Multimer structural prediction

The predicted complex structures, pLDDT plots, and PAE plots for MRGPRX4-RAMP2 were generated with AlphaFold Colab, which uses a slightly simplified version of AlphaFold v2.1.0^16, 17^. The endogenous signal sequence for RAMP2 was omitted from the input. The pdb file of each predicted complex was loaded into PDBePISA^66^ [https://www.ebi.ac.uk/msd-srv/prot_int/cgi-bin/piserver] to generate a list of interacting residues and calculate interaction surface area. Pairing of interacting residues and assignment of interaction type was done manually in ChimeraX^21, 22^. Image generation was carried out in PyMol (The PyMOL Molecular Graphics System, Version 2.0 Schrödinger, LLC).

## Data availability

The authors declare that all other data supporting the findings of this study are available within the paper and supplementary figures.

## Acknowledgements

We thank the Rockefeller University’s Fisher Drug Discovery Resource Center (DDRC) for advice and access to instrumentation (LI-COR Odyssey M (LI-COR), Biotek Synergy Neo, and BioTek Synergy NEO-TRF Hybrid multi-mode reader (BioTek Instruments). We thank Dr. Alison J. North, Dr. Tao Tong and Dr. Ved Sharma at the Rockefeller University’s Bio-imaging Resource Center (BIRC) for training, discussion, assistance with the experiments, and access to instrumentation (DeltaVision Image Restoration Inverted Olympus IX-71 microscope and Imaris software). We thank Marcus Saarinen, Leo Dahl and Annika Bendes for discussion. The PRESTO-Tango plasmid kit was a gift from Bryan Roth (Addgene kit # 1000000068). We thank the Rockefeller University Structural Biology Resource Center (SBRC; RRID: SCR_017732) for assistance with structural prediction analysis. We thank everyone at the Human Protein Atlas and the team of the Affinity Proteomics Unit in Stockholm. E. C. was funded by the François Wallace Monahan Fellowship and the Tom Haines Fellowship in Membrane Biology. I.B.K. was supported by the Tri-Institutional Training Program in Chemical Biology, the National Institute of Health Grant T32 [GM136640], the Nicholson Short-Term Exchange, the Alexander Mauro Fellowship, and the Robertson Therapeutic Development Fund with support from the Denise and Michael Kellen Foundation through the Kellen Women’s Entrepreneurship Fund. This work was supported by funds for the Human Protein Atlas (HPA) and the Wallenberg Center for Protein Research (WCPR) provided by the Knut and Alice Wallenberg Foundation. Financial support was also provided by the KTH Center for Applied Precision Medicine (KCAP), the Erling-Persson Family Foundation.

## Contributions

E.C. and I.B.K. designed experiments, conducted experiments, compiled data, and analyzed data presented in this study. E.C., I.B.K. and T.P.S. designed the concept of the project. E.L. designed and conducted experiments. K.K. conducted experiments. T.D.C. analyzed data. I.B.K. and D.A.O. performed *in silico* structural predictions and analyses. I.B.K, E.C. and T.P.S. wrote the manuscript. M.H.D. and T.H. provided resources for the development and optimization of the Tet-On constructs design and assays. T.P.S. and J.S. supervised and coordinated the project.

## Ethics Declaration

Competing interests: The authors declare no competing interests

## Notes

### Competing Interest Statement

The authors have declared no competing interest.

