## Supplementary for "Itch receptor MRGPRX4 interacts with the receptor activity-modifying proteins (RAMPs)"

**Supplementary Information**

**Supplementary Figures (pages 2 – 13)**

Supplementary Fig. 1, Chemical structures of selected MRGPRX4 agonists and IP1 dose-response curves for nateglinide and DCA as a function of MRGPRX4 expression level.
Supplementary Fig. 2, Validation of mAbs to detect engineered and solubilized RAMPs and MRGPRX4 via epitope tags.
Supplementary Fig. 3, The effect of RAMP co-expression on nateglinide and DCA dose-response curves.
Supplementary Fig. 4, Characterization of the signaling properties of the different MRGPRX4, CALCRL, and RAMP constructs used in this study.
Supplementary Fig. 5, Optimization of the Tet-On inducible expression system using Tet-On FLAG-NLuc-HT7-MRGPRX4-GFP10-1D4 construct.
Supplementary Fig. 6, Map of interacting residues for Alpha-Fold Multimer MRGPRX4-RAMP2 complex structure.

**Supplementary Tables (pages 14 – 21)**

Supplementary Table 1. Fitting parameters of IP1 Dose-response curves in Figure 1, Figure 3, Supplementary Fig. 4, and Supplementary Fig. 5.
Supplementary Table 2. Statical test parameters for figures assessed by ordinary one-way ANOVA followed by Dunnett's multiple comparisons test.
Supplementary Table 3. Two-Phase Decay Model for Time-courses best-fit values.
Supplementary Table 4. MRGPRX4-RAMP2 predicted interacting residues, based on the Alpha-Fold Multimer predicted structure and the PDBePISA interface tool. The interaction types are based on manual annotation with ChimeraX. Model confidence indicates the predicted local distance difference test (pLDDT) value for each residue.
Supplementary Table 5. All oligonucleotides used to generate the various constructs used in this paper. **5a** TagMaster site-directed mutagenesis primers used to swap the FLAG epitope tag with a 3xHA epitope tag in FLAG-RAMP-OLLAS constructs. **5b** NEBuilder HiFi DNA Assembly primers used to amplify fragments and build the GPCR constructs used in this study. All oligonucleotides were ordered from IDT at the standard desalting grade.

**Supplementary References (page 22)**

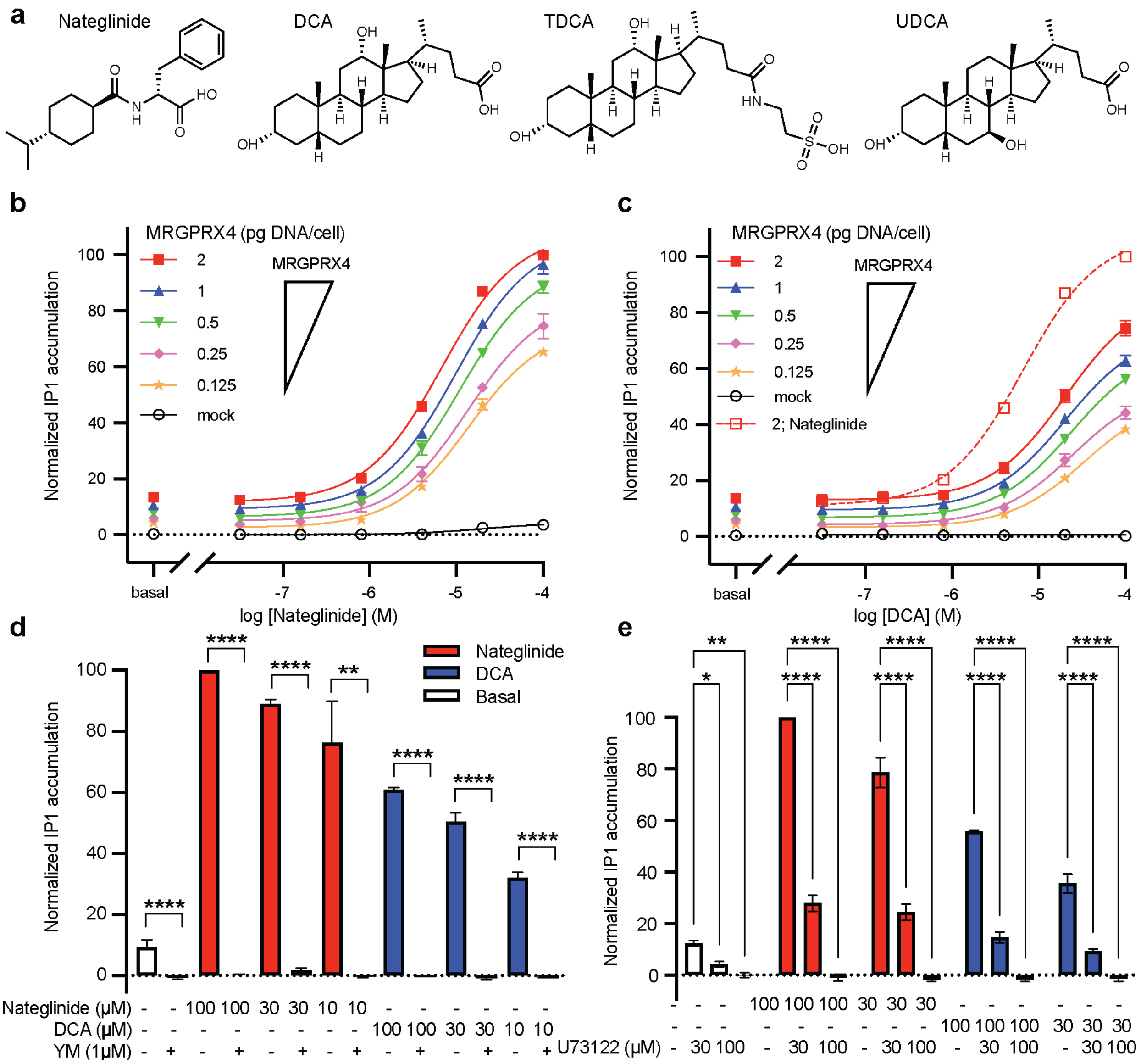
a
Nateglinide
DCA
TDCA
UDCA
b
MRGPRX4 (pg DNA/cell)
2
1
0.5
0.25
0.125
mock
MRGPRX4
Normalized IP1 accumulation
basal
log [Nateglinide] (M)
c
MRGPRX4 (pg DNA/cell)
2; Nateglinide
log [DCA] (M)
d
Nateglinide
DCA
Basal
Nateglinide (μM)
DCA (μM)
YM (1μM)
e
U73122 (μM)

**Supplementary Fig. 1, Chemical structures of selected MRGPRX4 agonists and IP1 dose-response curves for nateglinide and DCA as a function of MRGPRX4 expression level. a** Chemical structures for nateglinide, DCA, TDCA, and UDCA. Dose-dependent IP1 accumulation for nateglinide (**b**) and DCA (**c**) in HEK293T cells transfected with five different amounts of MRGPRX4-encoding plasmid (red square, 2pg/cell; blue triangle, 1pg/cell; green reverse triangle, 0.5pg/cell; pink diamond, 0.25pg/cell; orange star, 0.125pg/cell) compared with mock-transfected cells (black open circle, 2pg/cell). The curves are fitted to log[agonist] versus response with a three-parameters model. Fitting parameters are provided in **Supplementary Table 1**. Data are expressed as the mean ± SEM of the normalized IP1 accumulation to 100 μM nateglinide-induced cells transfected with 2pg/cell of MRGPRX4 plasmid as the maximum, and mock basal as the minimum. These results were used to prepare **Fig. 1 b** and **c.** Data are from three independent experiments performed in four technical replicates. **d, e** IP1 accumulation mediated by MRGPRX4 in the presence of nateglinide or DCA with or without the Gq inhibitor YM (**d**) or the phospholipase C (PLC) inhibitor U73122 (**e**). Data are expressed as the mean ± SEM of the normalized IP1 accumulation to 100 μM nateglinide and are from three (**d**) and two (**e**) independent experiments performed in four technical replicates. Statistical significance was determined by unpaired two-tailed t test (for data in panel **d**) or ordinary two-way ANOVA followed by Dunnett's multiple comparisons test (for data in panel **e**) (see **Supplementary Table 2)**. Statistical significance, ****$p < 0.0001$, **$p < 0.01$, * $p < 0.1$.

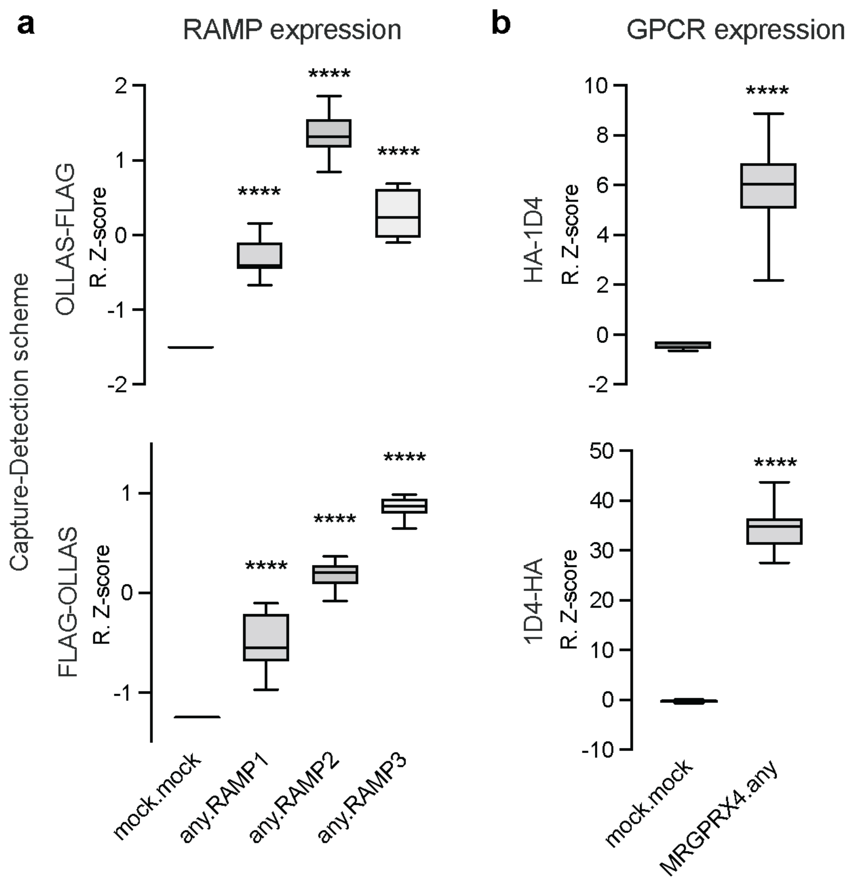
a
RAMP expression
b
GPCR expression
Capture-Detection scheme
OLLAS-FLAG
R. Z-score
FLAG-OLLAS
R. Z-score
HA-1D4
R. Z-score
1D4-HA
R. Z-score
****
mock.mock
any.RAMP1
any.RAMP2
any.RAMP3
mock.mock
MRGPRX4.any

**Supplementary Fig. 2, Validation of mAbs to detect engineered and solubilized RAMPs and MRGPRX4 via epitope tags. a** Freestyle293 cells were transfected with either epitope-tagged HA-MRGPRX4-1D4 or FLAG-RAMP-OLLAS, or co-transfected pairwise with MRGPRX4 and each RAMP. Lysates were incubated with the SBA, which included beads conjugated to each of the four mAbs targeting HA, 1D4, FLAG, and OLLAS. Each RAMP was captured with anti-OLLAS mAb beads and detected with PE-conjugated anti-FLAG mAb (top) and captured with anti-FLAG mAb beads and detected with PE-conjugated anti-OLLAS mAb (bottom). **b** MRGPRX4 was captured with anti-HA mAb beads and detected with PE-conjugated anti-1D4 mAb (top) and captured with anti-1D4 mAb beads and detected with PE-conjugated anti-HA mAb (bottom). Data are plotted as Robust Z-scores (R.Z-scores) and represent measurements from three independent experiments performed in duplicate. The box and whiskers plots represent the maximum and minimum extents of the measured values. Asterisks indicates that signal was significantly higher than mock.mock, as determined by ordinary one-way ANOVA followed by Dunnett's multiple comparisons test (for data in panel **a**) or unpaired two-tailed t test (for data in panel **b**) (see **Supplementary Table 2** for numerical parameters) (****$p < 0.0001$).

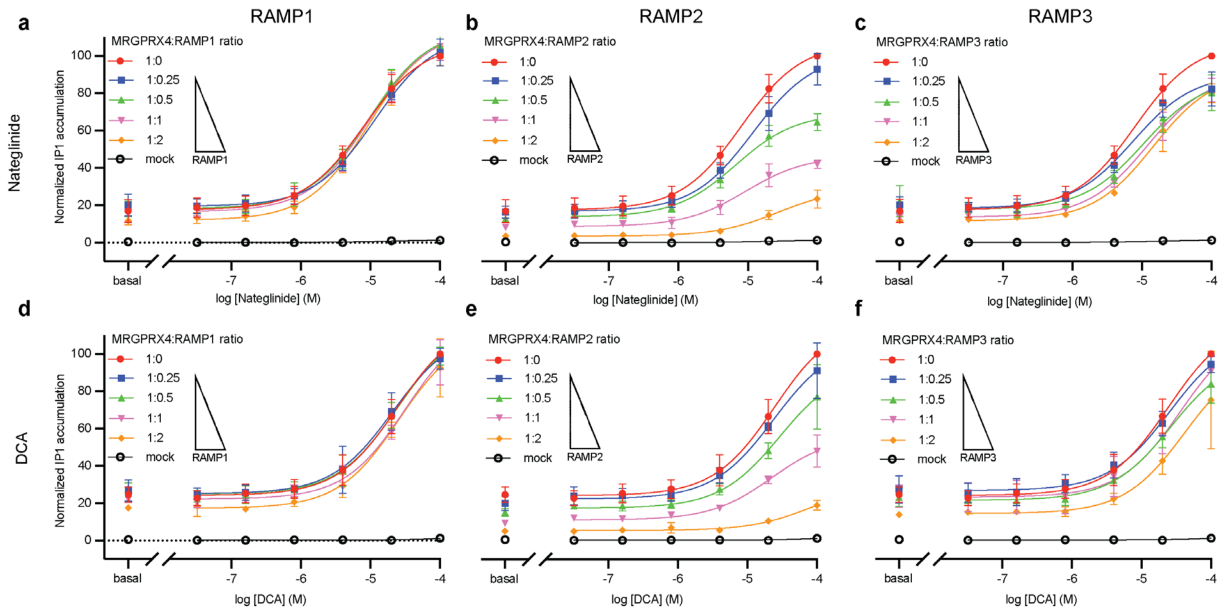
a
b
c
d
e
f
RAMP1
RAMP2
RAMP3
Nateglinide
DCA
Normalized IP1 accumulation
MRGPRX4:RAMP1 ratio
MRGPRX4:RAMP2 ratio
MRGPRX4:RAMP3 ratio
1:0
1:0.25
1:0.5
1:1
1:2
mock
basal
log [Nateglinide] (M)
log [DCA] (M)

**Supplementary Fig. 3, The effect of RAMP co-expression on nateglinide and DCA dose-response curves. a-c** IP1 accumulation was measured in response to nateglinide in cells expressing MRGPRX4 and increasing amounts of RAMP1 (**a**), RAMP2 (**b**) or RAMP3 (**c**). **d-f** IP1 accumulation was measured in response to DCA in cells expressing MRGPRX4 and increasing amounts of RAMP1 (**d**), RAMP2 (**e**) or RAMP3 (**f**). MRGPRX4:RAMP ratios are as follows: 1:0 red circle, 1:0.25 blue square, 1:0.5 green triangle, 1:1 pink reverse triangle, 1:2 orange diamond, and mock black open circle. These results were used to prepare **Fig. 4** in the main text. The curves are fits of the dose-response data to log[agonist] versus response with a three-parameters model. Fitting parameters are provided in **Supplementary Table 1**. Data are expressed as the mean ± SEM. Normalized IP1 accumulation is for each agonist separately. Data are from three independent experiments performed in four technical replicates.

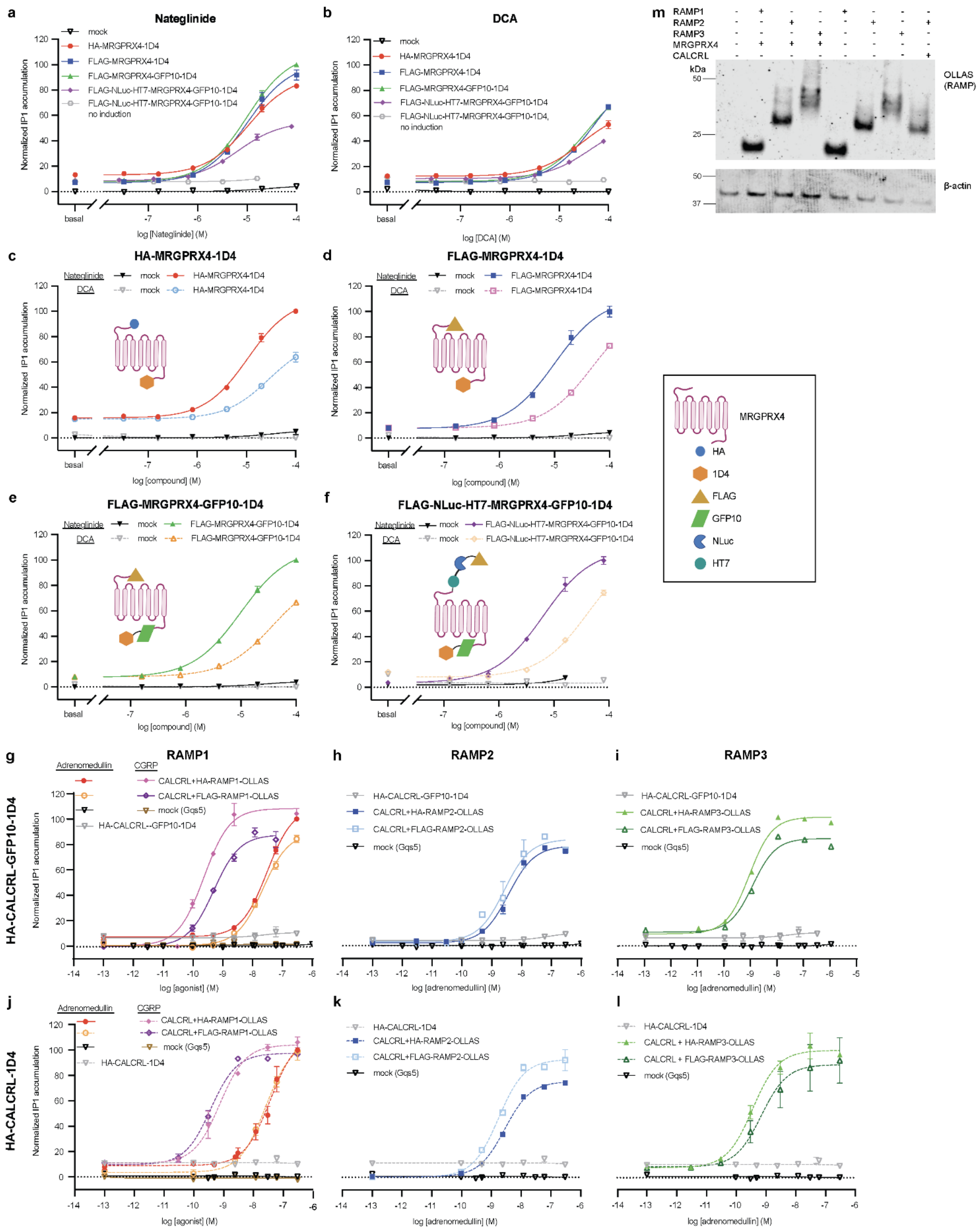
a
Nateglinide
mock
HA-MRGPRX4-1D4
FLAG-MRGPRX4-1D4
FLAG-MRGPRX4-GFP10-1D4
FLAG-NLuc-HT7-MRGPRX4-GFP10-1D4
FLAG-NLuc-HT7-MRGPRX4-GFP10-1D4 no induction
Normalized IP1 accumulation
basal
log [Nateglinide] (M)
b
DCA
mock
HA-MRGPRX4-1D4
FLAG-MRGPRX4-1D4
FLAG-MRGPRX4-GFP10-1D4
FLAG-NLuc-HT7-MRGPRX4-GFP10-1D4
FLAG-NLuc-HT7-MRGPRX4-GFP10-1D4, no induction
log [DCA] (M)
m
RAMP1
RAMP2
RAMP3
MRGPRX4
CALCRL
kDa
OLLAS (RAMP)
β-actin
c
HA-MRGPRX4-1D4
Nateglinide
DCA
mock
HA-MRGPRX4-1D4
log [compound] (M)
d
FLAG-MRGPRX4-1D4
FLAG-MRGPRX4-1D4
MRGPRX4
HA
1D4
FLAG
GFP10
NLuc
HT7
e
FLAG-MRGPRX4-GFP10-1D4
f
FLAG-NLuc-HT7-MRGPRX4-GFP10-1D4
g
RAMP1
Adrenomedullin
CGRP
CALCRL+HA-RAMP1-OLLAS
CALCRL+FLAG-RAMP1-OLLAS
mock (Gqs5)
HA-CALCRL--GFP10-1D4
HA-CALCRL-GFP10-1D4
log [agonist] (M)
h
RAMP2
HA-CALCRL-GFP10-1D4
CALCRL+HA-RAMP2-OLLAS
CALCRL+FLAG-RAMP2-OLLAS
mock (Gqs5)
log [adrenomedullin] (M)
i
RAMP3
HA-CALCRL-GFP10-1D4
CALCRL+HA-RAMP3-OLLAS
CALCRL+FLAG-RAMP3-OLLAS
mock (Gqs5)
j
HA-CALCRL-1D4
Adrenomedullin
CGRP
CALCRL+HA-RAMP1-OLLAS
CALCRL+FLAG-RAMP1-OLLAS
mock (Gqs5)
HA-CALCRL-1D4
k
HA-CALCRL-1D4
CALCRL+HA-RAMP2-OLLAS
CALCRL+FLAG-RAMP2-OLLAS
mock (Gqs5)
l
HA-CALCRL-1D4
CALCRL + HA-RAMP3-OLLAS
CALCRL + FLAG-RAMP3-OLLAS
mock (Gqs5)

**Supplementary Fig. 4, Characterization of the signaling properties of the different MRGPRX4, CALCRL, and RAMP constructs used in this study**.

**a-f** IP1 accumulation in cells expressing the different MRGPRX4 constructs used in the study. All dose-response curves are normalized to the maximum IP1 accumulation for nateglinide-stimulated FLAG-MRGPRX4-GFP10-1D4. **a, b** IP1 accumulation dose-responses of **a** nateglinide and **b** DCA mediated by HA-MRGPRX4-1D4 (red circle), FLAG-MRGPRX4-1D4 (blue square), FLAG-MRGPRX4-GFP10-1D4 (green triangle), and Tet-On FLAG-NLuc-HT7-MRGPRX4-GFP10-1D4 with dox (purple diamond) and without dox (open gray circle). **c-f** Dose-responses of nateglinide (filled symbol, solid line) and DCA (open symbol, dashed line) for each MRGPRX4 construct compared with mock. **c** HA-MRGPRX4-1D4 (used in **Fig.2, Supplementary Fig.2**), **d** FLAG-MRGPRX4-1D4, **e** FLAG-MRGPRX4-GFP10-1D4 (used in **Fig.1,3,4,5, Supplementary Fig. 1,3**), and **f** Tet-On FLAG-NLuc-HT7-MRGPRX4-GFP10-1D4 with dox compared with mock (no dox) (used in **Fig.6, Supplementary Fig. 5**). **g-l** IP1 accumulation in cells expressing the different RAMP constructs in the presence of an epitope-tagged CALCRL. Dose-responses of adrenomedullin (all RAMPs) and calcitonin gene-related peptide (CGRP) (RAMP1 only) promoted by HA-CALCRL-GFP10-1D4 (used in **Fig.5**) or HA-CALCRL-1D4 (used in the GPCR-RAMP interactome SBA pilot[1]) in complex with either HA-RAMP-OLLAS (used in **Fig.3,4,5,6, Supplementary Fig. 3, Supplementary Fig. 5**) or FLAG-RAMP-OLLAS (used in **Fig.2, Supplementary Fig. 2** and in the GPCR-RAMP interactome SBA pilot[1]). Cells transfected with CALCLR-RAMP2 served as the positive control for RAMP functionality. IP1 accumulation is normalized to adrenomedullin stimulated FLAG-CALCRL-GFP10-1D4 + RAMP1 (**g-i**) or HA-CALCRL-1D4 + RAMP1 (**j-l**). Data are expressed as the mean ± SEM of normalized IP1 accumulation and are from three independent experiments performed in technical triplicate. Fitting parameters are in **Supplementary Table 1**. **m** OLLAS Ab was used for immunodetection of each epitope-tagged RAMP construct (HA-RAMP-OLLAS) in the presence or absence of FLAG-MRGPRX4-GFP10-1D4 or HA-CALCRL-GFP10-1D4. (Uncropped immunoblots in **Source data**).

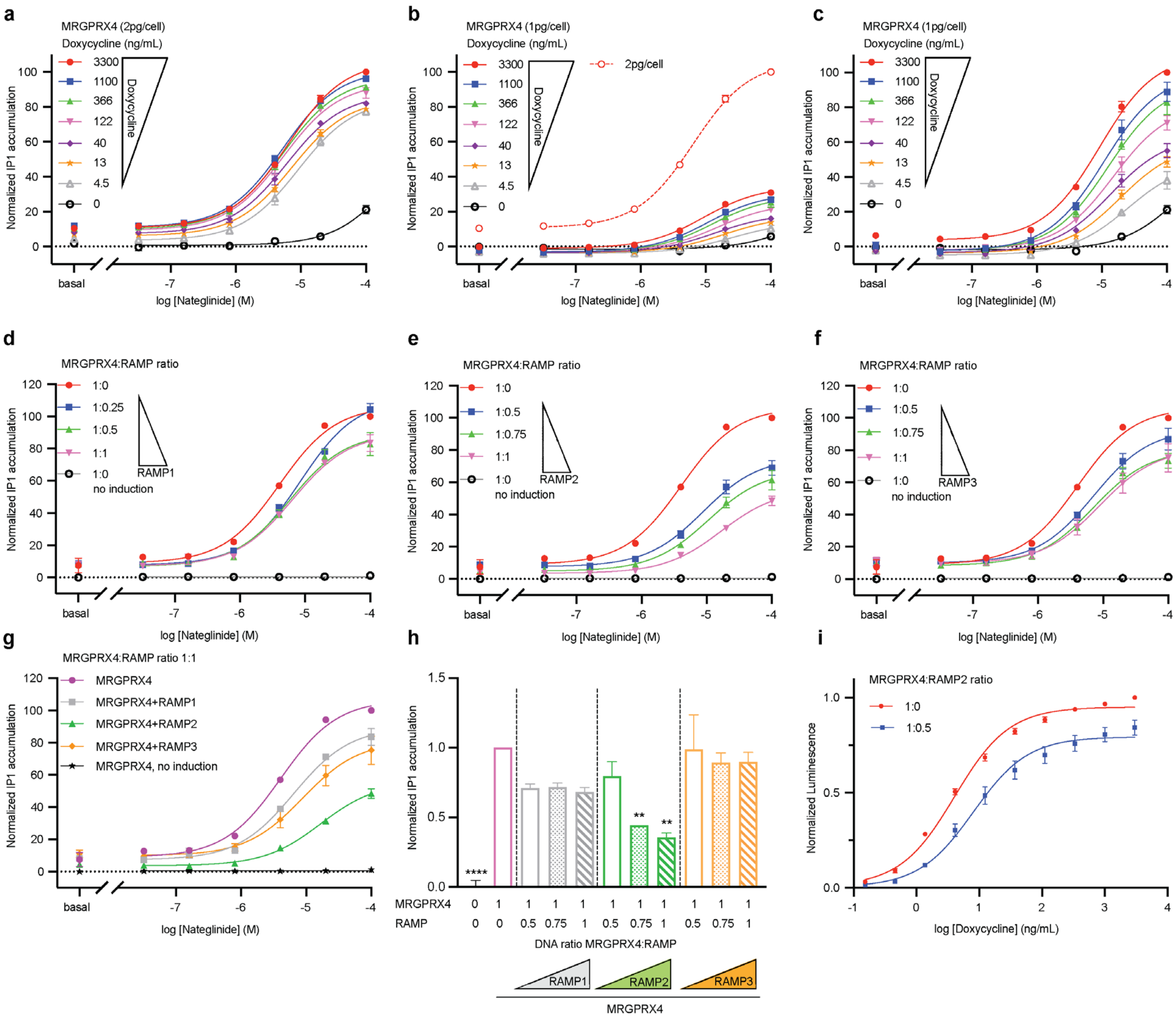

a
MRGPRX4 (2pg/cell)
Doxycycline (ng/mL)
3300
1100
366
122
40
13
4.5
0
Doxycycline
Normalized IP1 accumulation
basal
log [Nateglinide] (M)
b
MRGPRX4 (1pg/cell)
Doxycycline (ng/mL)
2pg/cell
c
MRGPRX4 (1pg/cell)
Doxycycline (ng/mL)
d
MRGPRX4:RAMP ratio
1:0
1:0.25
1:0.5
1:1
1:0
no induction
RAMP1
e
MRGPRX4:RAMP ratio
1:0
1:0.5
1:0.75
1:1
1:0
no induction
RAMP2
f
MRGPRX4:RAMP ratio
1:0
1:0.5
1:0.75
1:1
1:0
no induction
RAMP3
g
MRGPRX4:RAMP ratio 1:1
MRGPRX4
MRGPRX4+RAMP1
MRGPRX4+RAMP2
MRGPRX4+RAMP3
MRGPRX4, no induction
h
Normalized IP1 accumulation
****
**
**
MRGPRX4
RAMP
DNA ratio MRGPRX4:RAMP
RAMP1
RAMP2
RAMP3
MRGPRX4
i
MRGPRX4:RAMP2 ratio
1:0
1:0.5
Normalized Luminescence
log [Doxycycline] (ng/mL)

**Supplementary Fig. 5, Optimization of the Tet-On inducible expression system using Tet-On FLAG-NLuc-HT7-MRGPRX4-GFP10-1D4 construct.**
**a-c** Three-dimensional titration of Tet-On MRGPRX4 constructs comparing receptor expression, receptor induction (dox titration with seven concentrations, up to 3300 ng/mL), and nateglinide dose-response (six concentrations and basal). Dose-response curves are normalized to IP1 accumulation in nateglinide-stimulated MRGPRX4 expressing cells (**a, b** 2 pg DNA/cell. **c** 1 pg DNA/cell). Data represent three independent experiments with four technical replicate each. **d-h** Effect of increasing amounts of each RAMP co-expressed with Tet-On MRGPRX4 construct on nateglinide-dependent IP1 accumulation. **d** RAMP1; **e** RAMP2; **f** RAMP3. **g** Effect of Tet-On MRGPRX4-RAMP co-expression (DNA ratio 1:1) on nateglinide-dependent IP1 accumulation (data are from **d-f**). Data are normalized to nateglinide stimulated-MRGPRX4. The curves are fits of the dose-response data to log[agonist] versus response with a three-parameters model. Fitting parameters are provided in **Supplementary Table 1**. **h** Effect of increasing amounts of RAMP co-expression on basal IP1 accumulation promoted by the Tet-On MRGPRX4 construct. Data are normalized to MRGPRX4 basal and represent two independent experiments with four technical replicate each. Statistical significance was determined by ordinary one-way ANOVA followed by Dunnett's multiple comparisons test to MRGPRX4 alone (see **Supplementary Table 2**) (****$p < 0.0001$, **$p < 0.01$, if not marked then not significant). **i** Dox dose-dependent receptor total expression following induction of Tet-On MRGPRX4, expressed alone or with RAMP2 (DNA ratio 1:0.5). Data are plotted as total luminescence normalized to MRGPRX4 alone (maximal concentration of dox) and are from three independent experiments with three replicates each. Data are displayed as the mean ± SEM.

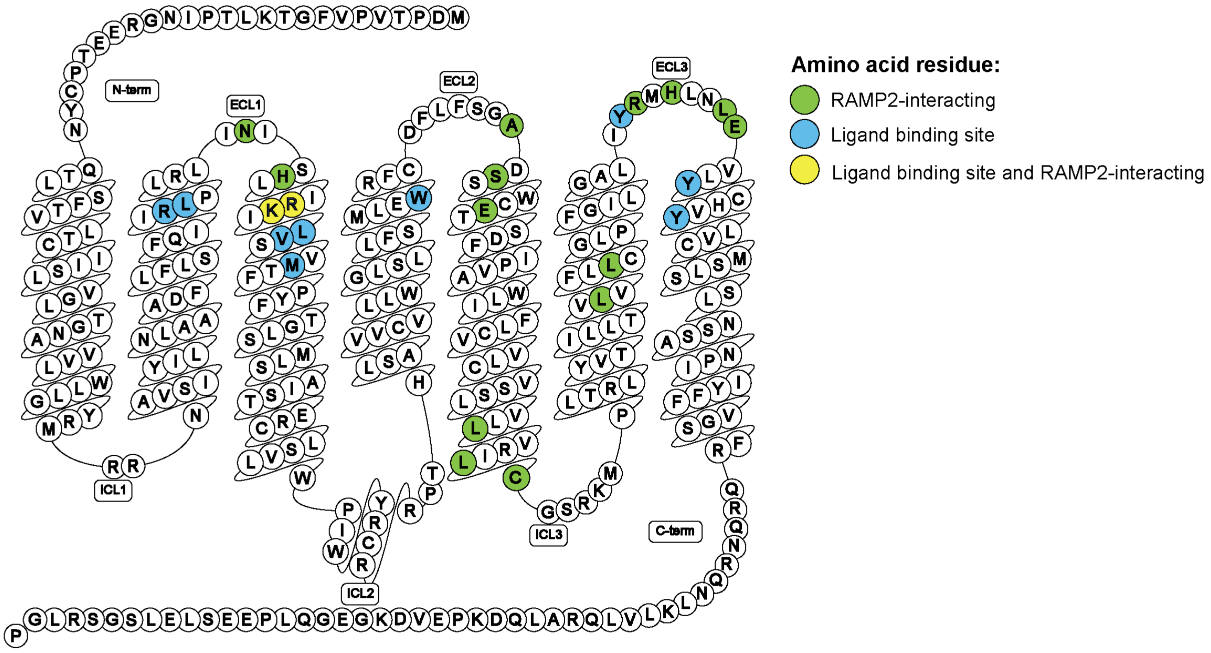
N-term
ECL1
ECL2
ECL3
ICL1
ICL2
ICL3
C-term
Amino acid residue:
RAMP2-interacting
Ligand binding site
Ligand binding site and RAMP2-interacting

**Supplementary Fig. 6, Map of interacting residues for Alpha-Fold Multimer MRGPRX4-RAMP2 complex structure.** Predicted complex formation between MRGPRX4 and RAMP2 was generated with Alpha-Fold Multimer Colab[2,3]. Snake plot diagram of MRGPRX4 showing the predicted RAMP2-interacting residues (green), ligand binding site residues (blue, for nateglinide analog MS47134[4]), and ligand-binding site residues that are also predicted to interact with RAMP2 (yellow). Plot generated with GPCRdb[5,6]. The full list of predicted interacting residues is provided in **Supplementary Table 4.**

**Supplementary Table 1**. Fitting parameters of IP1 Dose-response curves in **Fig. 1, Fig. 3, Supplementary Fig. 4,** and **Supplementary Fig. 5**.

| | Basal | $E_{max}$ | $logEC_{50}$ | $EC_{50}$ | span | Dof [a)] |
|---|---|---|---|---|---|---|
| MRGPRX4 (**Fig. 1a**) | | | | | | |
| Nateglinide | 14.95 | 102.6 | -4.973 | 1.065e-5 | 87.64 | 90 |
| DCA | 15.03 | 77.92 | -4.716 | 1.921e-5 | 62.89 | 85 |
| TDCA | 14.54 | 84.17 | -4.274 | 5.321e-5 | 69.63 | 77 |
| UDCA | 13.66 | 24956 | -0.9961 | 0.1009 | 24924 | 77 |
| MRGPRX4 (Dose-response Nateglinide) (**Fig.S1a**) | | | | | | |
| 2 pg | 11.77 | 108.0 | -5.168 | 6.797e-6 | 96.26 | 18 |
| 1 pg | 9.219 | 105.9 | -5.009 | 9.786e-6 | 96.66 | 18 |
| 0.5 pg | 6.421 | 97.45 | -4.958 | 1.102e-5 | 91.03 | 18 |
| 0.25 pg | 4.926 | 84.39 | -4.865 | 1.364e-5 | 79.46 | 18 |
| 0.125 pg | 2.563 | 74.80 | -4.856 | 1.392e-5 | 72.24 | 18 |
| MRGPRX4 (Dose-response DCA) (**Supplementary Fig. 1b**) | | | | | | |
| 2 pg | 12.99 | 87.15 | -4.692 | 2.031e-5 | 74.16 | 18 |
| 1 pg | 9.471 | 73.82 | -4.691 | 2.039e-5 | 64.35 | 18 |
| 0.5 pg | 6.783 | 68.02 | -4.622 | 2.389e-5 | 61.23 | 18 |
| 0.25 pg | 4.249 | 54.24 | -4.609 | 2.461e-5 | 49.99 | 18 |
| 0.125 pg | 3.308 | 50.32 | -4.470 | 3.390e-5 | 47.01 | 18 |
| MRGPRX4 + titration RAMP1 (Dose-response Nateglinide) (**Supplementary Fig. 3a**) | | | | | | |
| 2 pg (of RAMP1) | 11.98 | 114.4 | -5.028 | 9.375e-6 | 102.4 | 18 |
| 1 pg | 16.52 | 113.9 | -5.025 | 9.448e-6 | 97.37 | 18 |
| 0.5 pg | 18.21 | 114.8 | -5.019 | 9.564e-6 | 96.60 | 18 |
| 0.25 pg | 19.44 | 111.9 | -4.939 | 1.152e-5 | 92.43 | 18 |
| MRGPRX4 + titration RAMP2 (Dose-response Nateglinide) (**Supplementary Fig. 3b**) | | | | | | |
| 2 pg | 3.623 | 28.83 | -4.574 | 2.664e-5 | 25.21 | 18 |
| 1 pg | 8.724 | 46.54 | -5.032 | 9.299e-5 | 37.81 | 17 |
| 0.5 pg | 13.85 | 69.53 | -5.175 | 6.685e-5 | 55.69 | 17 |
| 0.25 pg | 16.83 | 101.3 | -4.928 | 1.181e-5 | 84.47 | 17 |
| MRGPRX4 + titration RAMP3 (Dose-response Nateglinide) (**Supplementary Fig. 3c**) | | | | | | |
| 2 pg | 11.70 | 91.35 | -4.839 | 1.450e-5 | 79.65 | 17 |
| 1 pg | 13.65 | 90.43 | -4.920 | 1.202e-5 | 76.78 | 17 |
| 0.5 pg | 17.78 | 88.29 | -4.977 | 1.055e-5 | 70.51 | 18 |
| 0.25 pg | 18.35 | 90.32 | -5.128 | 7.440e-6 | 71.98 | 17 |
| MRGPRX4 + titration RAMP1 (Dose-response DCA) (**Supplementary Fig. 3d**) | | | | | | |
| 2 pg | 17.30 | 111.8 | -4.597 | 2.532e-5 | 94.50 | 18 |
| 1 pg | 22.25 | 118.0 | -4.504 | 3.136e-5 | 95.79 | 18 |
| 0.5 pg | 24.94 | 117.0 | -4.615 | 2.424e-5 | 92.02 | 18 |
| 0.25 pg | 25.16 | 112.8 | -4.683 | 2.077e-5 | 87.65 | 18 |
| MRGPRX4 + titration RAMP2 (Dose-response DCA) (**Supplementary Fig. 3e**) | | | | | | |
| 2 pg | 5.412 | 29.86 | -4.097 | 8.007e-5 | 24.45 | 20 |
| 1 pg | 11.18 | 56.18 | -4.652 | 2.226e-5 | 44.99 | 20 |
| 0.5 pg | 17.50 | 94.46 | -4.530 | 2.952e-5 | 76.96 | 20 |
| 0.25 pg | 22.40 | 107.2 | -4.625 | 2.371e-5 | 84.83 | 20 |
| MRGPRX4 + titration RAMP3 (Dose-response DCA) (**Supplementary Fig. 3f**) | | | | | | |
| 2 pg | 14.52 | 100.4 | -4.384 | 4.128e-5 | 85.85 | 18 |
| 1 pg | 23.28 | 115.4 | -4.436 | 3.667e-5 | 92.07 | 18 |
| 0.5 pg | 21.60 | 100.4 | -4.574 | 2.664e-5 | 78.85 | 18 |
| 0.25 pg | 26.80 | 111.0 | -4.596 | 2.533e-5 | 84.19 | 18 |
| MRGPRX4 (compare constructs) (Dose-response Nateglinide) (**Supplementary Fig. S4a,c-f**) | | | | | | |
| HA-MRGPRX4-1D4 | 13.13 | 91.49 | -4.972 | 1.066e-5 | 78.36 | 18 |
| FLAG-MRGPRX4-1D4 | 6.903 | 102.0 | -4.994 | 1.015e-5 | 95.07 | 18 |
| FLAG-MRGPRX4-GFP10-1D4 | 7.722 | 109.7 | -5.004 | 9.908e-6 | 101.9 | 18 |
| FLAG-NLuc-HT7-MRGPRX4-GFP10-1D4 | 8.575 | 55.22 | -5.193 | 6.407e-6 | 46.65 | 18 |

| | | | | | | |
|---|---|---|---|---|---|---|
| MRGPRX4 (compare constructs) (Dose-response DCA) (**Supplementary Fig. 4b,c-f**) | | | | | | |
| Mock | 2.330 | 0.2364 | -7.834 | 1.465e-8 | -2.094 | 18 |
| HA-MRGPRX4-1D4 | 12.52 | 64.71 | -4.544 | 2.855e-5 | 52.19 | 18 |
| FLAG-MRGPRX4-1D4 | 7.469 | 96.26 | -4.308 | 4.920e-5 | 88.79 | 18 |
| FLAG-MRGPRX4-GFP10-1D4 | 8.165 | 88.54 | -4.424 | 3.764e-5 | 80.37 | 18 |
| FLAG-NLuc-HT7-MRGPRX4-GFP10-1D4 | 10.56 | 55.05 | -4.388 | 4.093e-5 | 44.49 | 18 |
| HA-CALCRL-GFP10-1D4 (Dose-response Adrenomedullin) (**Supplementary Fig.4g-i**) | | | | | | |
| Mock | 0.1993 | 11.54 | -5.176 | 6.665e-6 | 11.34 | 42 |
| CALCRL | 4.246 | 10.49 | -7.129 | 7.435e-8 | 6.243 | 11 |
| CALCRL+HA-RAMP1-OLLAS | 7.282 | 109.6 | -7.512 | 3.074e-008 | 102.4 | 17 |
| CALCRL+FLAG-RAMP1-OLLAS | 0.4611 | 89.41 | -7.655 | 2.213e-008 | 88.95 | 18 |
| CALCRL+HA-RAMP2-OLLAS | 2.278 | 78.81 | -8.436 | 3.665e-9 | 76.54 | 11 |
| CALCRL+FLAG-RAMP2-OLLAS | 3.344 | 83.87 | -8.581 | 2.626e-9 | 80.53 | 10 |
| CALCRL+HA-RAMP3-OLLAS | 9.590 | 101.6 | -9.089 | 8.142e-010 | 92.04 | 17 |
| CALCRL+FLAG-RAMP3-OLLAS | 11.09 | 84.86 | -8.955 | 1.109e-009 | 73.76 | 18 |
| HA-CALCRL-GFP10-1D4 (Dose-response CGRP) (**Supplementary Fig.4g**) | | | | | | |
| Mock | 0.4062 | 1.291 | -9.835 | 1.461e-10 | 0.8853 | 41 |
| CALCRL | 6.580 | 10.82 | -7.655 | 2.212e-8 | 4.236 | 18 |
| CALCRL+HA-RAMP1-OLLAS | 0.3849 | 105.5 | -9.635 | 2.317e-010 | 105.1 | 12 |
| CALCRL+FLAG-RAMP1-OLLAS | -0.3006 | 87.64 | -9.305 | 4.957e-010 | 87.94 | 9 |
| HA-CALCRL-1D4 (Dose-response Adrenomedullin) (**Supplementary Fig.4j-l**) | | | | | | |
| Mock | 0.9133 | 0.05397 | -7.376 | 4.210e-008 | -0.8593 | 30 |
| CALCRL | 11.35 | Unstable | -0.7361 | 0.1836 | Unstable | 30 |
| CALCRL+HA-RAMP1-OLLAS | 9.024 | 111.5 | -7.444 | 3.599e-008 | 102.5 | 30 |
| CALCRL+FLAG-RAMP1-OLLAS | 3.676 | 106.5 | -7.596 | 2.538e-008 | 102.9 | 27 |
| CALCRL+HA-RAMP2-OLLAS | 0.3834 | 75.53 | -8.549 | 2.827e-009 | 75.15 | 18 |
| CALCRL+FLAG-RAMP2-OLLAS | 0.1775 | 92.88 | -8.758 | 1.746e-009 | 92.70 | 17 |
| CALCRL+HA-RAMP3-OLLAS | 8.590 | 108.1 | -9.450 | 3.544e-010 | 99.49 | 9 |
| CALCRL+FLAG-RAMP3-OLLAS | 9.152 | 95.99 | -9.162 | 6.881e-010 | 86.83 | 11 |
| HA-CALCRL- 1D4 (Dose-response CGRP) (**Supplementary Fig.4j**) | | | | | | |
| Mock | Unstable | -0.8959 | -16.10 | 7.881e-017 | Unstable | 24 |
| CALCRL+HA-RAMP1-OLLAS | 10.86 | 104.1 | -9.130 | 7.412e-010 | 93.21 | 9 |
| CALCRL+FLAG-RAMP1-OLLAS | 9.681 | 97.50 | -9.435 | 3.673e-010 | 87.81 | 9 |
| FLAG-NLuc-HT7-MRGPRX4-GFP10-1D4 (Titration doxycycline) (2pg/cell) (**Supplementary Fig. 5a**) | | | | | | |
| Dox 3300 ng/mL | 10.81 | 106.4 | -5.197 | 6.360e-6 | 95.60 | 18 |
| Dox 1100 ng/mL | 11.03 | 101.4 | -5.267 | 5.302e-6 | 90.38 | 18 |
| Dox 366 ng/mL | 9.980 | 97.01 | -5.274 | 5.320e-6 | 87.03 | 18 |
| Dox 122 ng/mL | 9.245 | 94.00 | -5.268 | 5.400e-6 | 84.76 | 18 |
| Dox 40 ng/mL | 7.421 | 87.55 | -5.217 | 6.065e-6 | 80.13 | 18 |

| | | | | | | |
|---|---|---|---|---|---|---|
| Dox 13 ng/mL | 6.149 | 84.94 | -5.134 | 7.354e-6 | 78.79 | 18 |
| Dox 4.5 ng/mL | 3.549 | 84.60 | -5.040 | 9.121e-6 | 81.05 | 18 |
| No dox | 0.7169 | 72.26 | -3.602 | 0.0002500 | 71.55 | 18 |
| FLAG-NLuc-HT7-MRGPRX4-GFP10-1D4 (Titration doxycycline) (1pg/cell) (**Supplementary Fig. 5b**) | | | | | | |
| Dox 3300 ng/mL | -1.008 | 34.37 | -5.033 | 9.258e-6 | 35.37 | 18 |
| Dox 1100 ng/mL | -3.039 | 30.65 | -4.968 | 1.077e-5 | 33.69 | 18 |
| Dox 366 ng/mL | -3.000 | 29.08 | -4.893 | 1.279e-5 | 32.08 | 18 |
| Dox 122 ng/mL | -2.879 | 24.93 | -4.821 | 1.510e-5 | 27.81 | 18 |
| Dox 40 ng/mL | -3.380 | 18.76 | -4.862 | 1.373e-5 | 22.14 | 18 |
| Dox 13 ng/mL | -3.433 | 17.37 | -4.718 | 1.915e-5 | 20.80 | 18 |
| Dox 4.5 ng/mL | -3.809 | 14.10 | -4.590 | 2.568e-5 | 17.91 | 18 |
| No dox | -1.503 | 24.99 | -3.577 | 0.0002650 | 26.49 | 18 |
| FLAG-NLuc-HT7-MRGPRX4-GFP10-1D4 (Titration doxycycline) (over 1pg/cell Normalized) (**Supplementary Fig. 5c**) | | | | | | |
| Dox 3300 ng/mL | 3.818 | 110.7 | -5.036 | 9.197e-6 | 106.8 | 18 |
| Dox 1100 ng/mL | -2.284 | 99.95 | -4.969 | 1.073e-5 | 102.2 | 18 |
| Dox 366 ng/mL | -2.161 | 94.31 | -4.900 | 1.258e-5 | 96.48 | 18 |
| Dox 122 ng/mL | -1.806 | 82.37 | -4.824 | 1.499e-5 | 84.18 | 18 |
| Dox 40 ng/mL | -3.289 | 63.50 | -4.868 | 1.355e-5 | 66.79 | 18 |
| Dox 13 ng/mL | -3.443 | 59.23 | -4.721 | 1.901e-5 | 62.67 | 18 |
| Dox 4.5 ng/mL | -4.623 | 48.88 | -4.601 | 2.506e-5 | 53.50 | 18 |
| No dox | -1.726 | 55.58 | -3.825 | 0.0001497 | 57.31 | 18 |
| FLAG-NLuc-HT7-MRGPRX4-GFP10-1D4 + RAMP1 (Titration RAMP1) (**Supplementary Fig. 5d**) | | | | | | |
| Ratio 1:0 | 8.926 | 106.7 | -5.392 | 4.059e-006 | 97.74 | 11 |
| Ratio 1:0.5 | 7.772 | 111.3 | -5.074 | 8.425e-006 | 103.5 | 25 |
| Ratio 1:0.75 | 6.875 | 90.02 | -5.225 | 5.950e-006 | 83.14 | 25 |
| Ratio 1:1 | 7.220 | 89.56 | -5.192 | 6.430e-006 | 82.34 | 25 |
| Ratio 1:0, no dox | 0.2822 | 2.457 | -3.895 | 0.0001274 | 2.175 | 21 |
| FLAG-NLuc-HT7-MRGPRX4-GFP10-1D4 + RAMP2 (Titration RAMP2) (**Supplementary Fig. 5e**) | | | | | | |
| Ratio 1:0 | 8.926 | 106.7 | -5.392 | 4.059e-006 | 97.74 | 11 |
| Ratio 1:0.5 | 7.610 | 75.64 | -5.058 | 8.757e-006 | 68.03 | 25 |
| Ratio 1:0.75 | 4.862 | 67.94 | -4.977 | 1.053e-005 | 63.08 | 11 |
| Ratio 1:1 | 3.554 | 55.74 | -4.778 | 1.668e-005 | 52.19 | 25 |
| Ratio 1:0, no dox | 0.2822 | 2.457 | -3.895 | 0.0001274 | 2.175 | 21 |
| FLAG-NLuc-HT7-MRGPRX4-GFP10-1D4 + RAMP3 (Titration RAMP3) (**Supplementary Fig. 5f**) | | | | | | |
| Ratio 1:0 | 8.926 | 106.7 | -5.392 | 4.059e-006 | 97.74 | 11 |
| Ratio 1:0.5 | 10.08 | 93.43 | -5.144 | 7.170e-006 | 83.35 | 25 |
| Ratio 1:0.75 | 8.392 | 80.54 | -5.148 | 7.112e-006 | 72.15 | 24 |
| Ratio 1:1 | 9.885 | 81.18 | -5.056 | 8.795e-006 | 71.30 | 25 |
| Ratio 1:0, no dox | 0.2822 | 2.457 | -3.895 | 0.0001274 | 2.175 | 21 |
| FLAG-NLuc-HT7-MRGPRX4-GFP10-1D4 Basal IP1(Titration RAMPs) (**Supplementary Fig. 5h**) | | | | | | |
| Ratio 1:0 | 1±0.000 | NA | NA | NA | NA | NA |
| RAMP1 Ratio 1:0.5 | 0.709±0.029 | NA | NA | NA | NA | NA |
| RAMP1 Ratio 1:75 | 0.716±0.029 | NA | NA | NA | NA | NA |
| RAMP1 Ratio 1:1 | 0.683±0.032 | NA | NA | NA | NA | NA |
| RAMP2 Ratio 1:0.5 | 0.794±0.104 | NA | NA | NA | NA | NA |
| RAMP2 Ratio 1:75 | 0.444±0.0007 | NA | NA | NA | NA | NA |
| RAMP2 Ratio 1:1 | 0.356±0.031 | NA | NA | NA | NA | NA |
| RAMP3 Ratio 1:0.5 | 0.990±0.246 | NA | NA | NA | NA | NA |
| RAMP3 Ratio 1:75 | 0.893±0.069 | NA | NA | NA | NA | NA |
| RAMP3 Ratio 1:1 | 1.072±0.179 | NA | NA | NA | NA | NA |
| FLAG-NLuc-HT7-MRGPRX4-GFP10-1D4 total expression (Titration dox) (**Supplementary Fig. 5i**) | | | | | | |
| MRGPRX4 | 0.009273 | 0.9497 | 0.6070 | 4.046 | 0.9404 | 87 |
| MRGPRX4+RAMP2 | 0.003719 | 0.7930 | 0.9067 | 8.067 | 0.7893 | 87 |

a) Degrees of Freedom,

**Supplementary Table 2.** Statical test parameters for figures assessed by ordinary one-way ANOVA followed by Dunnett's multiple comparisons test or two-tailed t test.

| Figure | Panel | DoF | F | Condition used for comparison | Notes |
|---|---|---|---|---|---|
| 1 | b (top) | 20 | 157.8 | MRGPRX4 (2pg/cell), for each agonist separately | |
| | b (bottom) | 24 | 243.3 | | |
| S1 | d | 4 | 2.623 | Basal | +/- YM for each condition. Unpaired, two-tailed t test. t-value = 15.57 |
| | d | 4 | Infinity | Ng 100μM | Unpaired, two-tailed t test. t-value = 205.2 |
| | d | 4 | 4.143 | Ng 30μM | Unpaired, two-tailed t test. t-value = 59.00 |
| | d | 4 | 641.3 | Ng 10μM | Unpaired, two-tailed t test. t-value = 5.711 |
| | d | 4 | 1.496 | DCA 100μM | Unpaired, two-tailed t test. t-value = 65.85 |
| | d | 4 | 22.08 | DCA 30μM | Unpaired, two-tailed t test. t-value = 17.86 |
| | d | 4 | 8.605 | DCA 10μM | Unpaired, two-tailed t test. t-value = 16.87 |
| | e | 15 | 60.56, 823.1, 121.7 | No U73122 treatment | F values correspond to interaction, row factor, column factor |
| 2 | b (Left) | 22 | 269.1 | mock.mock (all) | Figure 2 Panel B LHS, top to bottom, 1D4 capture, FLAG detection |
| | b (Left) | 28 | 197.8 | | 1D4 capture, OLLAS detection |
| | b (Left) | 35 | 55.03 | | HA capture, FLAG detection |
| | b (Left) | 35 | 46.56 | | HA capture, OLLAS detection |
| | b (Right) | 36 | 143.4 | | Figure 2 Panel B RHS, top to bottom, FLAG capture, 1D4 detection |
| | b (Right) | 31 | 258.3 | | FLAG capture, HA detection |
| | b (Right) | 34 | 113.4 | | OLLAS capture, 1D4 detection |
| | b (Right) | 30 | 71.16 | | OLLAS capture, HA detection |
| S2 | a (top) | 35 | 123.5 | mock.mock (all) | OLLAS capture, FLAG detection |
| | a (bottom) | 38 | 204.8 | | FLAG capture, OLLAS detection |
| | b (top) | 24 | 124.8 | | HA capture, 1D4 detection. Unpaired, two-tailed t test. t-value = 8.689. |
| | b (bottom) | 24 | 203.9 | | 1D4 capture, HA detection. Unpaired, two-tailed t test. t-value = 19.42. |
| 3 | b | 55 | 25.79 | Mock | |
| 4 | a (top) | 51 | 83.69 | MRGPRX4 1:0 | nateglinide or DCA- stimulated |
| | a (bottom) | 54 | 26.24 | MRGPRX4 1:0 | MRGPRX4 1:0 |
| | b | 64 | 40.94 | MRGPRX4 1:0 | MRGPRX4 1:0 basal |
| | c | 14 | 82.92 | MRGPRX4 1:0 | MRGPRX4,1:0+nateglinide100μM |
| | d | 14 | 71.03 | MRGPRX4 1:0 | MRGPRX4,1:0+DCA100μM |

| | | | | | |
|---|---|---|---|---|---|
| | e | 14 | 83.63 | MRGPRX4 1:0 | MRGPRX4,1:0+TDCA100μM |
| | f | 14 | 57.25 | MRGPRX4 1:0 | MRGPRX4,1:0+UDCA100μM |
| 5 | d | 52 | 1.854 | Basal (all) | β-arrestin1 + MRGPRX4: basal versus 100μM nateglinide Unpaired, two-tailed t test. t-value = 2.313 |
| | d | 52 | 7.531 | | β-arrestin2 + MRGPRX4: basal versus 100μM nateglinide Unpaired, two-tailed t test. t-value = 26.04 |
| | d | 52 | 7.448 | | β-arrestin1 + CALCRL + RAMP2: basal versus 200nM adrenomedullin Unpaired, two-tailed t test. t-value = 14.25 |
| | d | 52 | 8.583 | | β-arrestin2 + CALCRL + RAMP2: basal versus 200nM adrenomedullin Unpaired, two-tailed t test. t-value = 15.59 |
| | f | 15 | 2.801 | | Basal |
| | f | 45 | 33.51 | | Nateglinide |
| | f | 15 | 4.4 | | DCA |
| 6 | b | 32 | 44.67 | MRGPRX4 alone (+dox) | |
| | c | 26 | 4.355 | MRGPRX4 alone | Normalized BRET Ratio (left axis) |
| | c | 26 | 5.792 | | Surface expression (right axis) |
| S5 | h | 22 | 9.979 | MRGPRX4 alone | |

**Supplementary Table 3.** Two-Phase Decay Model for Time-courses best-fit values.

| | $y_0$ | $x_0$ | Plateau | $K_{fast}$ | $K_{slow}$ | Dof a) | Half-life (slow) | Half-life (fast) | Tau (slow) | Tau (fast) | Rate constant ratio |
|---|---|---|---|---|---|---|---|---|---|---|---|
| MRGPRX4-GFP10 BRET$^2$ time-course (**Fig. 5**) | | | | | | | | | | | |
| 100 μM b) | 0.01291 | 56.70 | 0.004639 | 0.008519 | 0.002534 | 64 | 273.5 | 81.37 | 394.6 | 117.4 | 3.361 |
| CALCRL-GFP10 + RAMP2 BRET$^2$ time-course (**Fig. 5**) | | | | | | | | | | | |
| 200 nM c) | 0.02190 | -468.5 | -0.03173 | 0.005142 | 0.005142 | 41 | 134.8 | 134.8 | 194.5 | 194.5 | 1.000 |

a) Degrees of Freedom
b) Nateglinide
c) Adrenomedullin

**Supplementary Table 4.** MRGPRX4-RAMP2 predicted interacting residues, based on the Alpha-Fold Multimer predicted structure and PDBePISA interface tool[7] [https://www.ebi.ac.uk/msd-srv/prot_int/cgi-bin/piserver]. The interaction types are based on manual annotation with ChimeraX[8,9]. Model confidence indicates the predicted local distance difference test (pLDDT) value for each residue. pLDDT values ≥ 50 are highlighted light green.

**4a.** MRGPRX4-RAMP2 (**Fig. 7**, **Supplementary Fig. 7**)

| MRGPRX4 residue | MRGPRX4 residue location a) | RAMP2 residue | Interaction type | MRGPRX4 model confidence | RAMP2 model confidence |
|---|---|---|---|---|---|
| Extracellular Interactions | | | | | |
| ASN89 | ECL1 | ASP66 | H bond | 42.4 | 67.7 |
| ASN89 | ECL1 | LEU67 | H bond (main chain) | 42.4 | 71.1 |
| ASN89 | ECL1 | GLU63 | H bond (main chain) | 42.4 | 70.8 |
| HIS92 | $3.22^{TM3}$ | GLU63 | H bond | 50.1 | 70.8 |
| HIS92 | $3.22^{TM3}$ | ASP56 | H bond | 50.1 | 72.5 |
| HIS92 | $3.22^{TM3}$ | GLU59 | H bond | 50.1 | 73.7 |
| ARG95 | $3.25^{TM3}$ | GLU59 | Salt bridge | 63.6 | 73.7 |
| ARG95 | $3.25^{TM3}$ | GLU63 | Salt bridge | 63.6 | 70.8 |
| LYS96 | $3.26^{TM3}$ | ASP56 | Salt bridge | 64.7 | 72.5 |
| ALA168 | ECL2 | ARG49 | H bond | 47.9 | 68.6 |
| SER170 | $5.29^{TM5}$ | ARG49 | van der Waals | 52.3 | 68.6 |
| GLU174 | $5.33^{TM5}$ | TRP44 | H bond | 64.3 | 66.6 |
| GLU174 | $5.33^{TM5}$ | ALA45 | H bond (main chain) | 64.3 | 61.4 |
| ARG241 | ECL3 | TRP44 | H bond (main chain) | 49.1 | 66.6 |
| HIS243 | ECL3 | TRP44 | Clash | 37.1 | 66.6 |
| HIS243 | ECL3 | TYR51 | Edge to face | 37.1 | 74.6 |
| HIS243 | ECL3 | HIS82 | H bond | 37.1 | 71.8 |
| LEU246 | ECL3 | TYR51 | van der Waals | 40.9 | 74.6 |
| LEU246 | ECL3 | ARG55 | van der Waals | 40.9 | 76.4 |
| GLU247 | ECL3 | PRO70 | van der Waals | 41.6 | 62.9 |
| Transmembrane Interactions | | | | | |
| Leu199 | $5.58^{TM5}$ | Ile118 | van der Waals | 83.4 | 48.6 |
| Leu199 | $5.58^{TM5}$ | Val122 | van der Waals | 83.4 | 48.3 |
| Leu203 | $5.62^{TM5}$ | Leu121 | van der Waals | 83.4 | 52.6 |
| Leu203 | $5.62^{TM5}$ | Val122 | van der Waals | 76.5 | 48.3 |
| Leu203 | $5.62^{TM5}$ | Arg125 | H bond | 76.5 | 51.8 |
| Cys204 | $5.63^{TM5}$ | Arg125 | H bond | 74.2 | 51.8 |
| Leu223 | $6.42^{TM6}$ | Ile115 | van der Waals | 91.9 | 44.5 |
| Leu223 | $6.42^{TM6}$ | Ile119 | van der Waals | 91.9 | 47.4 |
| Leu223 | $6.42^{TM6}$ | Ile118 | van der Waals | 91.9 | 48.6 |
| Leu227 | $6.46^{TM6}$ | Ile112 | van der Waals | 90.6 | 45.4 |
| Leu227 | $6.46^{TM6}$ | Ile115 | van der Waals | 90.6 | 44.5 |

a) MRGPRX4 residues are numbered according to the Ballesteros–Weinstein numbering obtained from GPCRdb.

**Supplementary Table 5. All oligonucleotides used to generate the various constructs used in this paper. 5a** TagMaster site-directed mutagenesis primers used to swap the FLAG epitope tag with a 3xHA epitope tag in FLAG-RAMP-OLLAS constructs. **5b** NEBuilder HiFi DNA Assembly primers used to amplify fragments and build the GPCR constructs used in this study. All oligonucleotides were ordered from IDT at the standard desalting grade.

**5a.** Primers for replacement of FLAG epitope tag with 3x HA tag for RAMPs

| Purpose of Primer | Oligonucleotide Sequence of Primers |
|---|---|
| RAMP1 Forward | CTG TTT ATG ACC ACC GCC TAC CCA TAC |
| RAMP1 Reverse | TAG TTG GCT TCC TGC AAT TGG CAA GCG |
| RAMP2 Forward | CCT CAC GAG GCC CTG GCC TAC CCA TA |
| RAMP2 Reverse | AGG CAG GGG CTG CAA TTG AGC GTA AT |
| RAMP3 Forward | TGT CCT AGA GCC GGC GGA TAC CCA TA |
| RAMP3 Reverse | CCT GTC TCG TTG CAC AAT TGA GCG TAA |

**5b.** NEBuilder HiFi DNA Assembly primers used in building constructs as described in Methods. Primers are grouped by construct generated (lines within table).

| Direction | Sequence of Primers (5' to 3') w/ overlaps underlined | Purpose |
|---|---|---|
| Forward (SP-FLAG-MRGPRX4) | CGT TTA AAC TTA AGC TTA GCG CCA CCA TGA AGA CGA TCA TC | Used in building FLAG-MRGPRX4-1D4 from PRESTO-Tango MRGPRX4 for PLA assay (1D4-pcDNA 3.1(+) vector was subject to enzymatic double digestion and purification). |
| Reverse (SP-FLAG-MRGPRX4) | TGC TGG CCT CAT CGA ATT CAC CGG TGC GTC CAC CGG TAT C | |
| Forward (SP-FLAG-MRGPRX4) | CTG CAC GAG ATG ATA CCG CCG CCA CCA TGA AGA CGA TC | Used in building FLAG-MRGPRX4-GFP10-1D4 for BRET[2] and $IP_1$ assays |
| Reverse (SP-FLAG-MRGPRX4) | CTC ACG AAT TCA CCG GTA CCG AAT TCA CCG GTG CGT CC | |
| Forward (GFP10-1D4 pcDNA 3.1(+) vector) | GGT ACC GGT GAA TTC GTG | |
| Reverse (GFP10-1D4 pcDNA 3.1(+) vector) | GGC GGT ATC ATC TCG TGC | |
| Forward (SP-FLAG-CALCRL) | CTG CAC GAG ATG ATA CCG CCG CCA CCA TGC GGC TGT GC | Used in building SP-FLAG-CALCRL-GFP10-1D4 |

| | | |
|---|---|---|
| Reverse (SP-FLAG-CALCRL) | CTC ACG AAT TCA CCG GTA CCG GTA CCG TTG TAC AGA TTC TCG GG | Note: Forward and Reverse (GFP10-1D4 pcDNA 3.1(+) vector) primers as the same as above |
| Forward (MRGPRX4-GFP10) | CCC CTG CAG GAG ATG ACA CCA TGG ATC CTA CTG TGC CTG TG | Used in building Tet-On SP-FLAG-Nluc-HT7-MRGPRX4-GFP10-1D4 for surface expression assays |
| Reverse (MRGPRX4-GFP10) | ACG GTG GTG CTG GCC TCA TCG GAT CCG CCT GCA GGC TT | |
| Forward (SP-FLAG-Nluc-HT7) | TTT GTA CAA AAA AGC AGG CTC CCA AGC TGG CTA GCG TTT AAA C | |
| Reverse (SP-FLAG-Nluc-HT7) | GGT GTC ATC TCC TGC AGG | |
| Forward (1D4 Tet-On vector) | GAT GAG GCC AGC ACC ACC | |
| Reverse (1D4 Tet-On vector) | AGC CTG CTT TTT TGT ACA AAC TTG C | |
| Forward (CALCRL-GFP10) | CCC CTG CAG GAG ATG ACA CCG AGC TGG AAG AGA GCC CC | Used in building Tet-On SP-FLAG-Nluc-HT7-CALCRL-GFP10-1D4 for surface expression assays |
| Reverse (CALCRL-GFP10) | ACG GTG GTG CTG GCC TCA TCC TTG TAC AGC TCG TCC ATG C | Note: Forward and Reverse (SP-FLAG-Nluc-HT7) and Forward and Reverse (1D4 Tet-On vector) primers are the same as above |
